# Conserved signals orchestrate self-organization and symmetry breaking of bi-layered epithelia during development and regeneration

**DOI:** 10.1101/2024.07.21.603898

**Authors:** Robin P. Journot, Mathilde Huyghe, Alexandre Barthelemy, Hugo Couto-Moreira, Jakub Sumbal, Marisa M. Faraldo, Maxime Dubail, Charles Fouillade, Silvia Fre

## Abstract

Organ development relies on complex molecular mechanisms that guide initially homogeneous populations of stem cells to differentiate into specialized cell types within defined spatial patterns. While stable during homeostasis, the proper spatial organization of cell types must be re-established in case of tissue injury for successful regeneration of organ shape and function. How cells commit to a differentiation path is a central question in stem cell research; however, the coordination between tissue geometry and cell fate specification remains enigmatic. To elucidate the molecular mechanisms instructing self-organization and symmetry breaking of epithelial stem cells, we developed a multi-faceted approach combining *in vitro* organoids, *ex vivo* embryonic tissue explants, and single-cell quantitative imaging to investigate the dynamic acquisition of cell fate in four bi-layered epithelia, during embryonic development but also in regeneration. Our findings indicate that tissue architecture is the primary determinant of cell fate decisions in these tissues. Upon the initial cell internalization event, the homogeneous population of stem cell break symmetry. Through genetic and pharmacological perturbations, we have demonstrated that a tightly coordinated interplay between Hippo/YAP and Notch signaling is essential for conveying information from tissue architecture to functional cell differentiation and stem cell potency restriction. Globally, this study uncovers the inherent capacity of stem cells to self-organize into multicellular structures, where the precise position of each differentiated cell is critical to instruct their differentiation choices during embryonic development and regeneration.

## Introduction

The mammary gland (MG), lacrimal gland (LG), salivary gland (SG), and prostate are all branched exocrine glands with a bi-layered epithelial architecture. These four organs originate from highly proliferative embryonic placodes composed of multipotent stem cells, derived from the ectoderm germ layer (for the MG, LG and SG), or from the endodermal urogenital sinus (UGS) for the prostate (Joseph et al., 2021; Pispa and Thesleff, 2003). During embryonic development, the homogenous epithelial placodes invaginate, form an epithelial bud that sinks further into the underlying mesenchyme and later ramify into a complex branched network.

The morphogenetic events that shape these tissues temporally coincide with the progressive potency restriction of initially multipotent stem cells into unipotent progenitors that sustain separate lineages of basal cells (BCs) and luminal cells (LCs) throughout further life (Basova et al., 2020; Centonze et al., 2020; Chatzeli et al., 2023; Choi et al., 2012; Davis et al., 2016; Lilja et al., 2018; May et al., 2018; Ousset et al., 2012; Tika et al., 2019; Van Keymeulen et al., 2017, 2011; Wang et al., 2017; Wuidart et al., 2018, 2016). The mature epithelium in the four organs is similarly organized, with BCs located in direct contact with the basement membrane (BM) and LCs facing the lumen. The epithelial cell types of the four organs are characterized by some conserved markers: p63 and the cytokeratins 5 (K5) and 14 (K14) mark BCs, while the cytokeratins 8 (K8) and 18 (K18) distinguish LCs (Delcroix et al., 2023; Sekiguchi et al., 2020; Toivanen and Shen, 2017; Watson and Khaled, 2020).

Perturbation of tissue homeostasis and organization in the MG, LG, SG and prostate lead to reactivation of multipotency in otherwise lineage-restricted stem cells, essential for proper tissue regeneration (Aure et al., 2015; Emmerson et al., 2018; Stingl et al., 2006; Wang et al., 2009). While both BCs and LCs participate in tissue regeneration, BCs exhibit a higher regenerative capacity (Centonze et al., 2020; Choi et al., 2012; Jamieson et al., 2017; Jardé et al., 2016; Kwak et al., 2016; Leong et al., 2008; Lu et al., 2013; Shackleton et al., 2006; Van Keymeulen et al., 2011; Wang et al., 2013; Yasuhara et al., 2022) compared to luminal cells (Guo et al., 2020; Karthaus et al., 2020; Rodilla et al., 2015; Wang et al., 2009; Weng et al., 2018).

While the processes dictating MG, LG, SG and prostate placode specification and epithelial branching have been thoroughly studied (Aa et al., 2003; Andl et al., 2002; Chatzeli et al., 2017; Z. Chen et al., 2014; Chu et al., 2004; Dean et al., 2005; Häärä et al., 2011; Mailleux et al., 2002; May et al., 2016; Mehta et al., 2013; Patel et al., 2011; Simons et al., 2012; van Genderen et al., 1994; Veltmaat et al., 2003, 2004, 2006; Wang et al., 2008), the molecular and mechanical underpinnings that instruct stem cell commitment have not been elucidated.

The Notch pathway is a master regulator of cell fate decisions, with a conserved role across species and in many tissues, such as the intestinal epithelium, skeletal muscle, epidermis, and pancreas (Baghdadi et al., 2018; Fre et al., 2005; Lloyd-Lewis et al., 2019; Seymour et al., 2020, 2020; Totaro et al., 2017). We have previously demonstrated that ectopic activation of Notch signaling in embryonic multipotent MG stem cells can skew their binary fate decision toward the luminal fate (Lilja et al., 2018) and recent reports suggest that Notch could be involved in the control of the lineage segregation in LG, SG and prostate (Chatzeli et al., 2023; Dvoriantchikova et al., 2017; Kuony and Michon, 2017; Wang et al., 2006; Wu et al., 2011).

The Hippo/YAP pathway has been reported to crosstalk with Notch signaling in several developmental and regenerative contexts (Totaro et al., 2018) and plays an essential role in tissue regeneration and stem cell self-renewal (Gregorieff et al., 2015; Loforese et al., 2017; Moya and Halder, 2019; von Gise et al., 2012; Yui et al., 2018). Ectopic activation of the YAP/TAZ pathway during homeostasis reactivates stem cell programs of self-renewal and multipotency, that are similar to the ones used for tissue regeneration (Panciera et al., 2016; Schlegelmilch et al., 2011; Zhao et al., 2014).

While YAP was shown to be functionally dispensable for the homeostasis of the virgin MG (Q. Chen et al., 2014), transient activation of YAP or inactivation of *Lats1/2* in adult MG LCs led to reactivation of the expression of some basal markers, notably K14 (Kern et al., 2022; Panciera et al., 2016). In the adult prostate, YAP knock-out displayed a weak phenotype, but genetic deletion during embryonic branching morphogenesis or prostate regeneration upon castration drastically affected both processes (Xie et al., 2023). Similarly, in the developing SG, YAP deletion resulted in defects in branching morphogenesis, while its forced activation promoted the expansion of cells in the salivary ductal compartment (Szymaniak et al., 2017).

Here, to elucidate the molecular mechanisms driving symmetry breaking and fate commitment in bi-layered epithelia, we combined *in vitro* organoids, *ex vivo* embryonic tissue explants, and single-cell quantitative imaging during mouse embryonic development and tissue regeneration. Such a multi-faceted approach allowed us to reveal the conserved temporal sequence of events by which multipotent stem cells self-organize and break their initial symmetry. Our findings indicate that positional cues act upstream of YAP patterning to regulate the precise timing of stem cell differentiation, which is in turn mediated by Notch lateral inhibition. The striking conservation of this signaling axis in four tissues derived from different embryonic germ layers, as well as the reactivation of similar processes during regeneration, demonstrate the robustness and precision of the elegant mechanism we unraveled.

## Results

### Dissociated adult BCs reactivate multipotency to give rise to self-organizing organoids recapitulating binary cell fate decisions of multipotent epithelial stem cells

To unravel the molecular mechanisms that determine the fate of embryonic epithelial stem cells, we developed an *in vitro* organoid culture system derived from single adult basal cells from four different tissues: the prostate, mammary gland (MG), lacrimal gland (LG), and salivary gland (SG). To this aim, we took advantage of the previously described capacity of adult mammary BCs to reactivate multipotency upon dissociation (Jamieson et al., 2017; Shackleton et al., 2006; Stingl et al., 2006) and tested if BCs isolated from the other three tissues also displayed such plasticity when grown as 3D organoids (Figure 1A). To document the multipotency of dissociated cells, we used a tamoxifen-inducible Cre recombinase specifically expressed in BCs thanks to the K5 promoter (*K5-Cre^ERT2^*) (Van Keymeulen et al., 2011), in combination with the *R26:mTmG* double fluorescent reporter line (Muzumdar et al., 2007) (*K5-Cre^ERT2^:mTmG*), allowing us to label *in vivo* adult BCs with membrane-tagged green fluorescent protein (mGFP) prior to cell isolation (Figure 1B). After isolation, the GFP-labelled BCs from each tissue were FACS-sorted using the established CD49f and EPCAM markers (Crowell et al., 2019; Stingl et al., 2006; Yasuhara et al., 2022) and seeded as single cells in Matrigel (Figures 1C-E, Supplementary Figure 1A-H). Cells were grown in tissue-specific organoid media (Crowell et al., 2019; Jardé et al., 2016; Kim et al., 2021) and formed clonal organoids consisting of hundreds of cells in a few days. We then examined cell fate composition of organoids using established differentiation markers for the two major cell types, K5 for BCs and K8 for LCs. After 7 days in culture, GFP^+^ organoids consisted of two spatially separated cell types: BCs (K5^+^) located at the periphery, and LCs (K8^+^) occupying the organoid inner part (Figure 1F). Importantly, we probed the organoid-forming capacity of single LCs as well and observed a much lower efficiency in organoid formation and no reactivation of multipotency (Supplementary figure 1I, J). We also investigated the expression of other classical cytokeratin markers of BCs and LCs, and confirmed the proper segregation of K14 to the external cells and of K18 to internal cells (Supplementary figure 1K).

**Figure 1.**
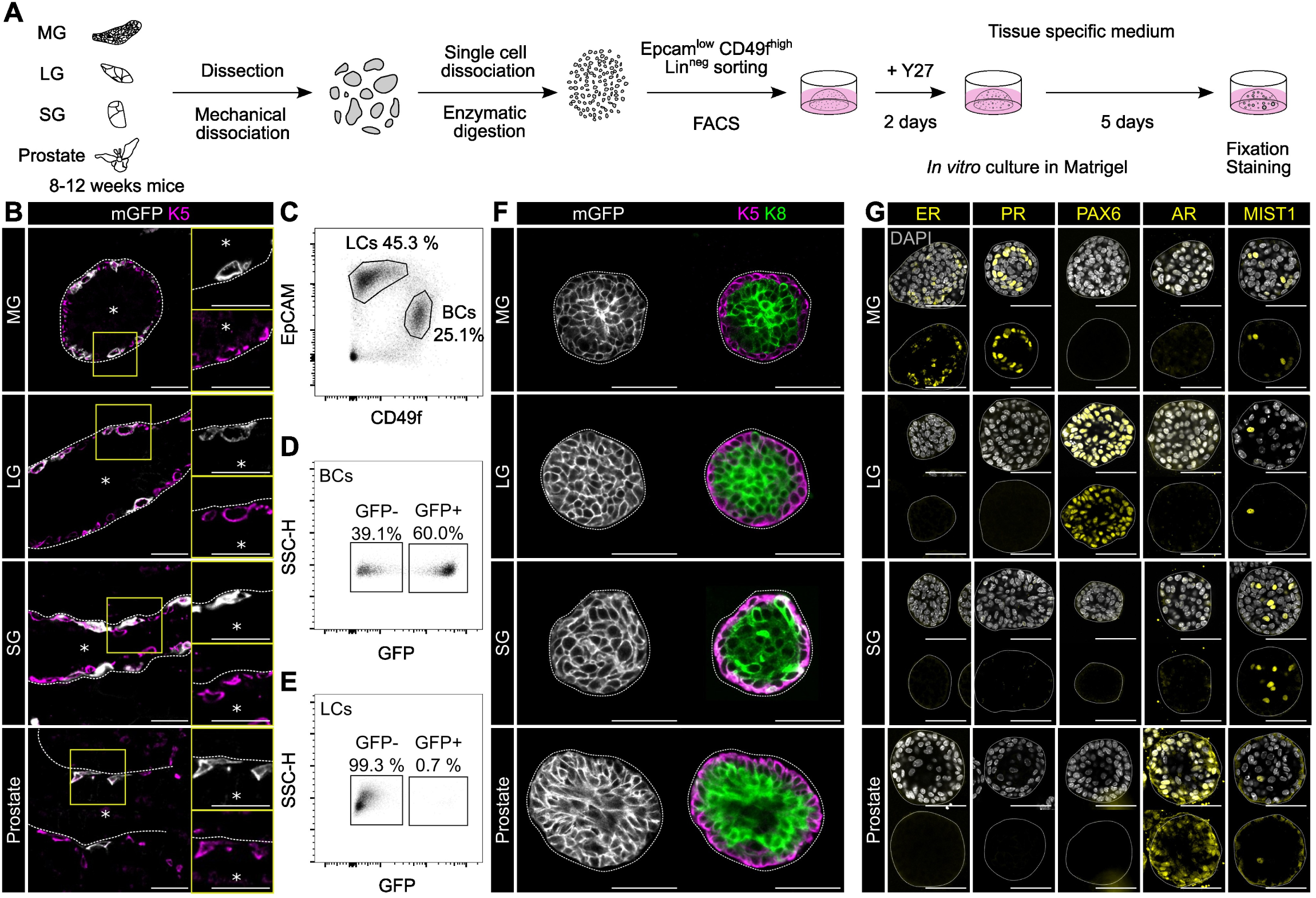
Dissociated adult BCs reactivate multipotency in organoids and recapitulate cell fate decisions of multipotent epithelial stem cells. **A**: Workflow of organoid development process from FACS sorted basal cells from adult MG, LG, SG and Prostate. **B:** Confocal imaging of native mGFP and immunostaining for K5 in *K5-Cre^ERT2^:mTmG* adult tissue section 72 h after intraperitoneal injection of TAM Scale bar: 50 µm. **C-E**: Sorting strategy to sort basal cells in MG. Suspension of single cells isolated from MG of adult *K5-Cre^ERT2^:mTmG* females 72 h after intraperitoneal injection of TAM, stained for non-epithelial markers (CD31 (APC), endothelium; CD45 (APC), immune cells; Ter119 (APC), erythroid lineage, i.e., Lin^pos^ cells), EpCAM (PE-Cy7) and CD49f (APC-Cy7) were gated to eliminate debris, doublets, Lin^pos^ and dead cells, positive to DAPI incorporation. EpCAM and CD49f expression was analized in alive Lin^neg^ cells. EpCAM^high^CD49^flow^ population correspond to LCs while the EpCAM^low^CD49f^high^ gate corresponds to BC population (C). GFP^+^ cells were selected in BCs and LCs populations (D, E). **F:** Confocal imaging of immunostaining for K5, K8 and native mGFP in organoids derived from GFP^+^ BCs. Scale bar: 50 µm. **G:** Confocal imaging of immunostaining for tissue markers in organoids derived from WT MG, LG, SG and Prostate after 7 days in culture. Scale bar: 50 µm.

To benchmark the organoids, we then selected a set of markers characteristic of each tissue of origin (Figure 1G). We confirmed that the mammary-specific markers (Estrogen receptor alpha, ERα and Progesterone receptor, PR), the prostate-specific marker Androgen Receptor (AR) and the lacrimal gland-specific transcription factor PAX6 were exclusively expressed in organoids derived from the corresponding tissue of origin. We further assessed the expression of a common marker of salivary, lacrimal and mammary cell types, MIST1, and confirmed that it was indeed expressed in SG, LG and MG-derived organoids.

Through these experiments we demonstrate the unique ability of adult BCs to reactivate multipotency in order to restore a luminal cell layer in all four tissues. Furthermore, we show that our organoid culture system faithfully recapitulates the binary cell fate decision of multipotent stem cells toward basal or luminal identity and reflects the correct tissue organization with external BCs and internal LCs, while retaining tissue-specific features. We thus used this minimal experimental system to study the mechanisms controlling the dynamics of fate segregation in stem-like cells.

### Fate segregation in single cell-derived organoids correlates with cell number

To characterize the dynamic events regulating binary fate decisions and assess their conservation in the four different tissues, we first examined the kinetics of lineage segregation. We fixed organoids at different time points and performed immunostaining for differentiation markers (Figure 2A). We observed an initial co-expression of the basal and luminal markers K5, K14 and K8 after 48 hours of culture, whereas lineage markers started to segregate radially after 72 hours in culture (Figure 2A, B, Supplementary Figure 2A). The vast majority of organoids derived from all four tissues presented an external layer of BCs and an inner mass of LCs by 144 hours of culture (Figure 2A, B). While fate segregation in organoids was asynchronous, we noticed that the organoid size critically distinguished fate-segregated (large) from non-segregated (small) organoids (Supplementary Figure 2B). To quantitatively determine the relationship between cell number and differentiation status, we designed a method to predict the number of cells in an organoid. All organoids displayed approximately a spherical shape across the whole time in culture, allowing us to use the area of the central optical section as a proxy for the total number of cells in each organoid (Figure 2C). We then investigated the dynamics of segregation of K5 and K8 expression in organoids by combining immunostaining with high-throughput imaging and single cell segmentation using computer vision algorithms to quantify K5 and K8 levels in individual cells (Figure 2D). We classified cells of organoids as “Internal” or “External” based on their distance to the edge of the organoids and computed the log ratio of the mean K8 and K5 intensities (Figure 2E, Supplementary figure 2C, See Methods). In small organoids (non-segregated fates), we observed a balanced expression of K5 and K8, resulting in a log-ratio around zero. In larger organoids, however, two cell populations became apparent, whose position was correlated with higher K8 levels (internal cells) or elevated K5 expression (external cells) (Figure 2E).

**Figure 2.**
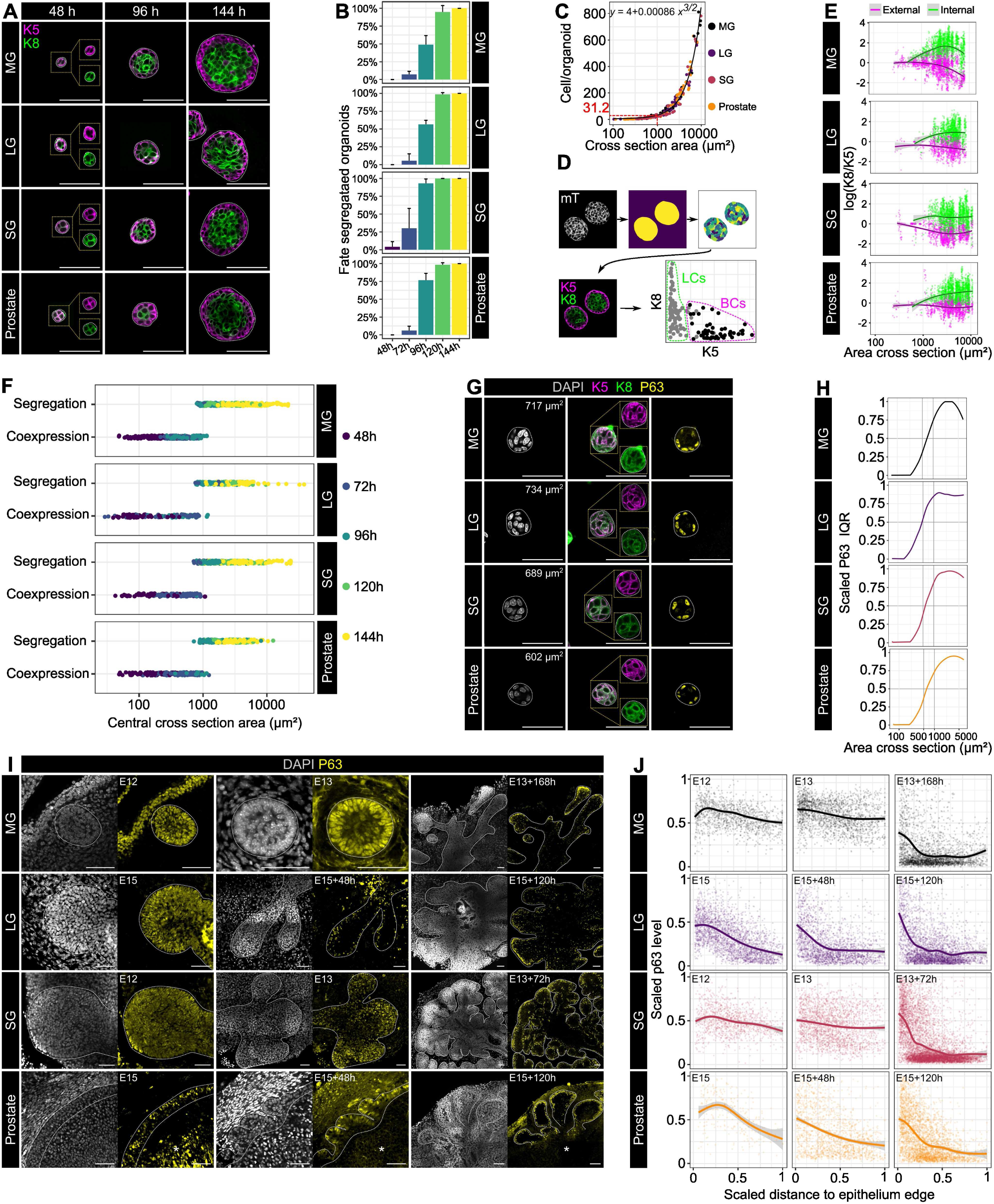
Fate segregation correlates with cell number. **A:** Confocal imaging of immunostaining for K5 and K8 in WT organoids at different stage of growth. Scale bar: 50 µm. **B:** Quantification of percentage of fate segregated organoids at different stage of growth. An organoid was classified as “fate segregated” when at least one K5^pos^K8^neg^ cell was identified. Data are displayed as Mean + s.d (n=3 independent cell sorting experiments). **C:** Relation between the cross-section area and the total number of cells contained in organoids of different size. **D:** Workflow of 2D organoid quantitative image analysis. Organoids are segmented based on Otsu’s thresholding and cells segmented using Cellpose on the membrane tdTomato channel. K5 or K8 intensities were then quantified within each mask. See Methods section for details of image analysis. **E:** K5 and K8 log ratio in cells from organoids of different size (n=3 independent cell sorting experiments). Cells were automatically classified as “Internal” or “External” based on the distance of the nucleus to the external boundary of the organoid. See Methods section for details of image analysis. **F:** Relation between the fate status and the central cross section area in organoids at different stage of growth (n=3 independent cell sorting experiments). **G:** Confocal imaging of immunostaining for K5, K8 and P63 in WT organoids after 96 hours in culture. Scale bar: 50 µm. **H:** Scaled interquartile range (Q3-Q1) of the P63 intensity measured in each cell of a center optical cross section of organoids of different size (n=3 independent cell sorting experiments). See Methods section. **I:** Confocal imaging of immunostaining for P63 in WT embryonic tissues at different stages of growth. Scale bar: 50 µm. **J:** Scaled of p63 intensities relative to the distance of the cell from the BM across development stages of embryonic MG, LG, SG and Prostate (n=3 biological replicates).

We thus compared the status of K5 and K8 segregation for individual organoids of different size and observed that segregation of K5 and K8 expression occurred in all four tissues around a sharp area threshold of ∼1000 µm² (Figure 2F), corresponding to 31 cells (Figure 2C). It is noteworthy that the tight and highly conserved correlation between the number of cells and fate segregation within an organoid explained the presence of a minority of small organoids in which cells still co-expressed K5 and K8 after 96 and 120 hours.

We next studied the early events leading to lineage segregation. *In vivo*, early embryonic stem cells composing all four epithelia express *Trp63* (gene encoding p63), which is progressively restricted to the basal lineage as development proceeds. Given the critical role of p63 in BCs specification and maintenance (Melino et al., 2015; Wuidart et al., 2018), we investigated the dynamics of p63 expression in organoids, using our image analysis pipeline based on organoid and nuclei segmentation (Supplementary figure 2C). Initially homogeneously expressed (Supplementary figure 2D), p63 became segregated to the external cells before K5 (Figure 2G). We precisely quantified the timing of p63 segregation by measuring the dispersion of p63 intensities within organoids of different size. The symmetry breaking event was defined as the moment at which the interquartile range, a measure of p63 expression heterogeneity, increased and reached half of its maximum value, which was between 500 and 750 µm² for p63, depending on the tissue (Figure 2H). The results of these experiments demonstrated that the size of the organoid is a reliable indicator of symmetry breaking. Accordingly, this quantifiable parameter was employed as a pseudotime to arrange organoids from several experiments along a continuous variable, representing the progression in cell differentiation.

### p63 progressively segregates to BCs during embryonic development of bi-layered epithelia

To assess the *in vivo* relevance of our findings, we performed the same measurements to evaluate the dynamics of fate markers segregation during embryonic development of the four bilayered epithelia. For this, we cultured embryonic tissue explants where the epithelial bud, initially composed of multipotent stem cells, is surrounded by its endogenous mesenchyme. This experimental setting faithfully recapitulates branching morphogenesis and fate commitment of the four epithelia (Supplementary figure 2E) (Berman et al., 2004; Carabaña et al., 2024; Kuony and Michon, 2017; Wang et al., 2021). We then assessed the levels of p63 at different timepoints and relative to the position of the epithelial cells within branching embryonic explants. In this *ex vivo* system, we could confirm the progressive downregulation of p63 in internal cells (Figure 2I-J), thereby demonstrating the conservation of the symmetry breaking of p63 expression in organoids and in *ex vivo* grown embryonic tissues.

### Notch signaling is necessary and sufficient for luminal fate determination in bilayered epithelia

To unravel the common molecular signals underlying fate commitment, we analyzed published scRNA-seq datasets of the four epithelial tissues under investigation during their development. We compared datasets from E15.5 MG (Carabaña et al., 2024), E16 LG (Farmer et al., 2017), E14 SG (Hauser et al., 2020) and E17.5 urogenital-sinus (the embryonic precursor of the prostate) (Lee et al., 2021), and identified a core set of basal and luminal markers shared among the four tissues (Supplementary figure 3A-N). Based on our previous work showing that Notch activation dictates luminal fate in any MG cell (Lilja et al., 2018), we decided to examinate further Notch signaling as a candidate to regulate the expression of common luminal markers.

To investigate the role of Notch in fate specification in all four tissues, we derived organoids from mice harboring the Cre-inducible, constitutively active intracellular domain of the Notch1 receptor alongside a nuclear GFP marker (*N1ICD-nGFP*) (Murtaugh et al., 2003) or only membrane-bound GFP (mGFP) as a control. We induced Cre-mediated recombination in *K5-Cre^ERT2^:N1ICD-nGFP* or *K5-Cre^ERT2^:mTmG* organoids by 4-hydroxytamoxifen (4-OHT) during the first 48h of organoid culture and analyzed cell type marker expression after 7 days of culture. While control mGFP^+^ cells had a K8^+^/K5^+^ cells ratio of approximately 2:1, N1ICD-nGFP^+^ mutant cells were invariably exclusively luminal (K8^+^) (Figure 3-A, B).

**Figure 3.**
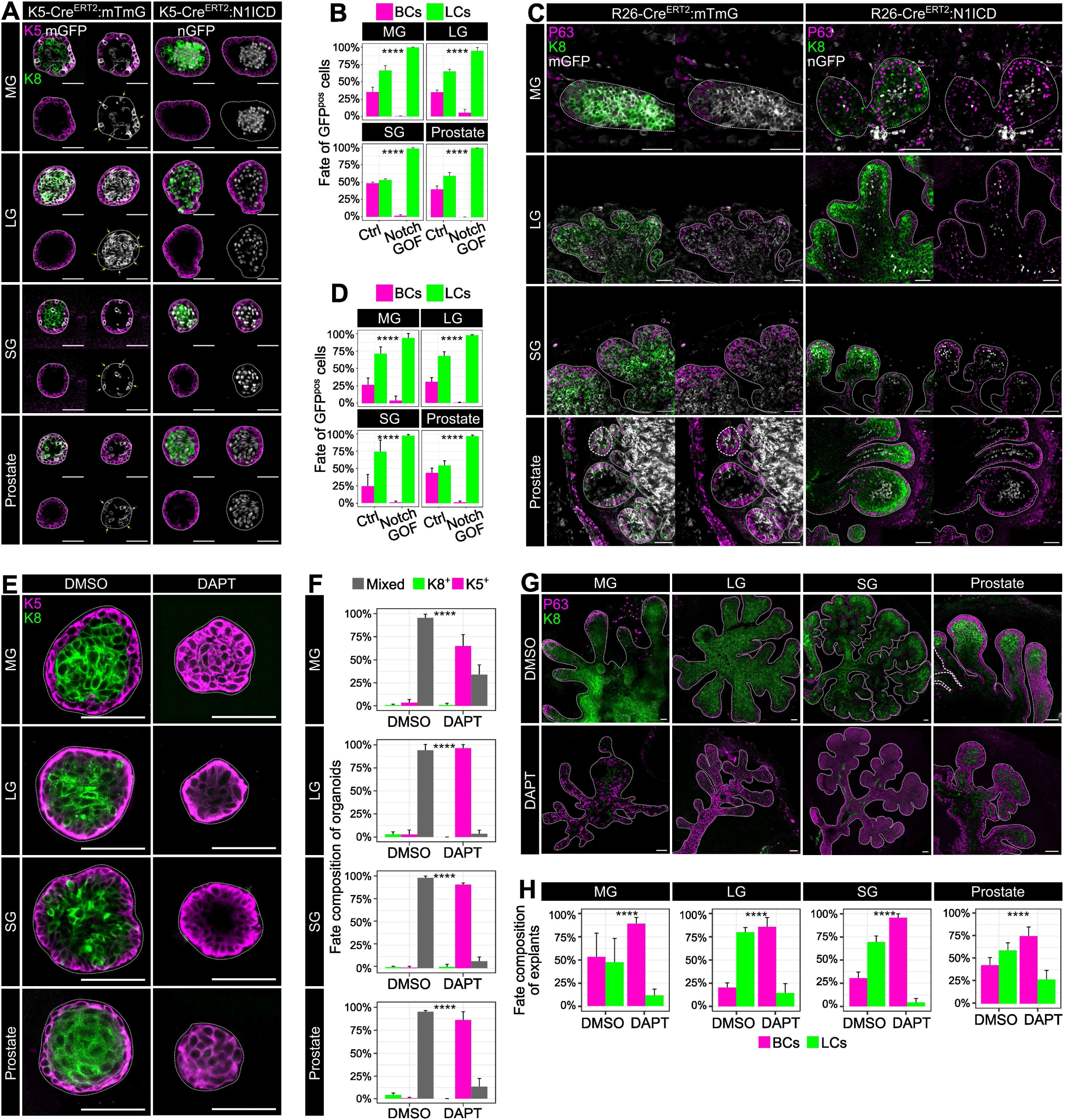
Notch signaling dictates cell fate decisions. **A**: Confocal imaging of immunostaining for K5, K8 and nGFP or native mGFP in *K5-Cre^ERT2^:mTmG* (Control) *of K5-Cre^ERT2^:N1ICD-nGFP* (Notch GOF) organoids respectively. Scale bar: 50 µm. **B:** Quantification of fate of mGFP^+^ (Control) or nGFP^+^ cells (Notch GOF). Data are displayed as Mean + s.d. Chi-squared test (n=3 independent cell sorting experiments). **C**: Confocal imaging of immunostaining for P63, K8 and GFP in *Rosa26-Cre^ERT2^:mTmG* (Control) or *Rosa26-Cre^ERT2^:N1ICD-nGFP* explants (Notch GOF). Scale bar: 50 µm. **D:** Quantification of fate of mGFP^+^ (Control) or nGFP^+^ cells (Notch GOF) in explants. Data are displayed as Mean + s.d. Chi-squared test (n=3 biological replicates). **E**: Confocal imaging of immunostaining for K5 and K8 in WT organoids treated for 7 days with DMSO or 50 µM DAPT. Scale bar: 50 µm. **F:** Quantification of percentage of organoids composed of K5^+^ and K8^+^ cells (Mixed), only BCs (K5^+^) or only luminal cells (K8^+^) in each condition. Data are displayed as Mean + s.d. Chi-squared test (n=3 independent cell sorting experiments). **G**: Confocal imaging of immunostaining for P63 and K8 in WT explants treated with DMSO or 50 µM DAPT for the whole duration of the culture. Scale bar: 50 µm. **H:** Quantification of fate of explants cells. Data are displayed as Mean + s.d. Chi-squared test (n=3 biological replicates).

We then corroborated these results by lineage tracing in embryonic tissues. Given that the K5 promoter was not efficient in targeting stem cells *in vivo* in all four embryonic tissues, we employed the inducible *R26-Cre^ERT2^*mouse line (Ventura et al., 2007) to target embryonic stem cells of the MG, SG, LG and prostate. Pregnant females harboring *R26-Cre^ERT2^:N1ICD-nGFP* or control *R26-Cre^ERT2^:mTmG* embryos were injected with tamoxifen at E12 to target MG and SG or at E14 to target LG and prostate cells. To prevent developmental abnormalities induced by ubiquitous Notch activation, we collected the embryos 24h later and cultured the embryonic tissues *ex vivo* as explants. The fate of mGFP^+^ control and N1ICD-nGFP^+^ mutant cells was then investigated by wholemount immunostaining (Figure 3C), when the epithelial explants had already branched and invaded the surrounding stroma (MG: E13+7d, SG: E13+3d, LG: E15+5d, Prostate: E15+5d). These experiments validated our results in organoids and established that N1ICD-expressing stem cells are invariably forced into a luminal fate, in all tissues investigated (Figure 3-D).

To assess if Notch signaling was indispensable for luminal differentiation, we then pharmacologically inhibited Notch activity in organoids using DAPT, a selective inhibitor of γ- secretase, the enzyme responsible for cleavage and activation of the Notch receptors (Geling et al., 2002). Organoids treated with DAPT for 7 days showed a striking increase in the percentage of K5^+^ BCs, with the vast majority of DAPT-treated organoids being exclusively composed of BCs (Figure 3E, F). To test the effect of Notch inhibition on embryonic multipotent stem cells, we cultivated explants derived from wild-type embryos with DAPT and quantified the cell type composition of the derived tissues. Consistent with our results in organoids, DAPT-treated embryonic explants were largely composed of BCs, unlike control explants presenting a physiological BCs to LCs ratio (Figure 3G, H). These experiments demonstrated that Notch signalling acts as the gatekeeper of luminal fate in all these tissues, being both necessary and sufficient for determining cell fate commitment of MG, SG, LG and prostate stem cells.

### Notch breaks symmetry following a radial pattern in both organoids and embryonic tissues

The discovery of an essential role for Notch signaling in the symmetry breaking of multipotent stem cells in the four tissues we examined prompted us to evaluate the dynamics of Notch pathway activation during organoid growth, and we decided to employ our *Hes1-emGFP* reporter mice (Fre et al., 2011). We first confirmed that expression of GFP from the *Hes*1 promoter faithfully mirrored the expression of the HES1 protein, a direct transcriptional target of Notch signaling (Jarriault et al., 1998) (Supplementary figure 4A). We then examined *Hes1* expression *in vivo* in adult MG, SG, LG and prostate and detected *Hes1-emGFP* exclusively in LCs in all four adult tissues (Supplementary Figure 4B). Next, we derived organoids from *Hes1-emGFP* mice and assessed the dynamics of *Hes1* expression. Whereas adult BCs *in vivo* never express Hes1-emGFP, cytoplasmic emGFP was rapidly detectable after seeding dissociated BCs and appeared homogeneously distributed in all reactivated cells for the first few rounds of cell divisions (Figure 4A). Upon symmetry breaking, however, *Hes1* expression became localized to internal cells and remained high after their commitment to LCs (Figure 4A).

**Figure 4.**
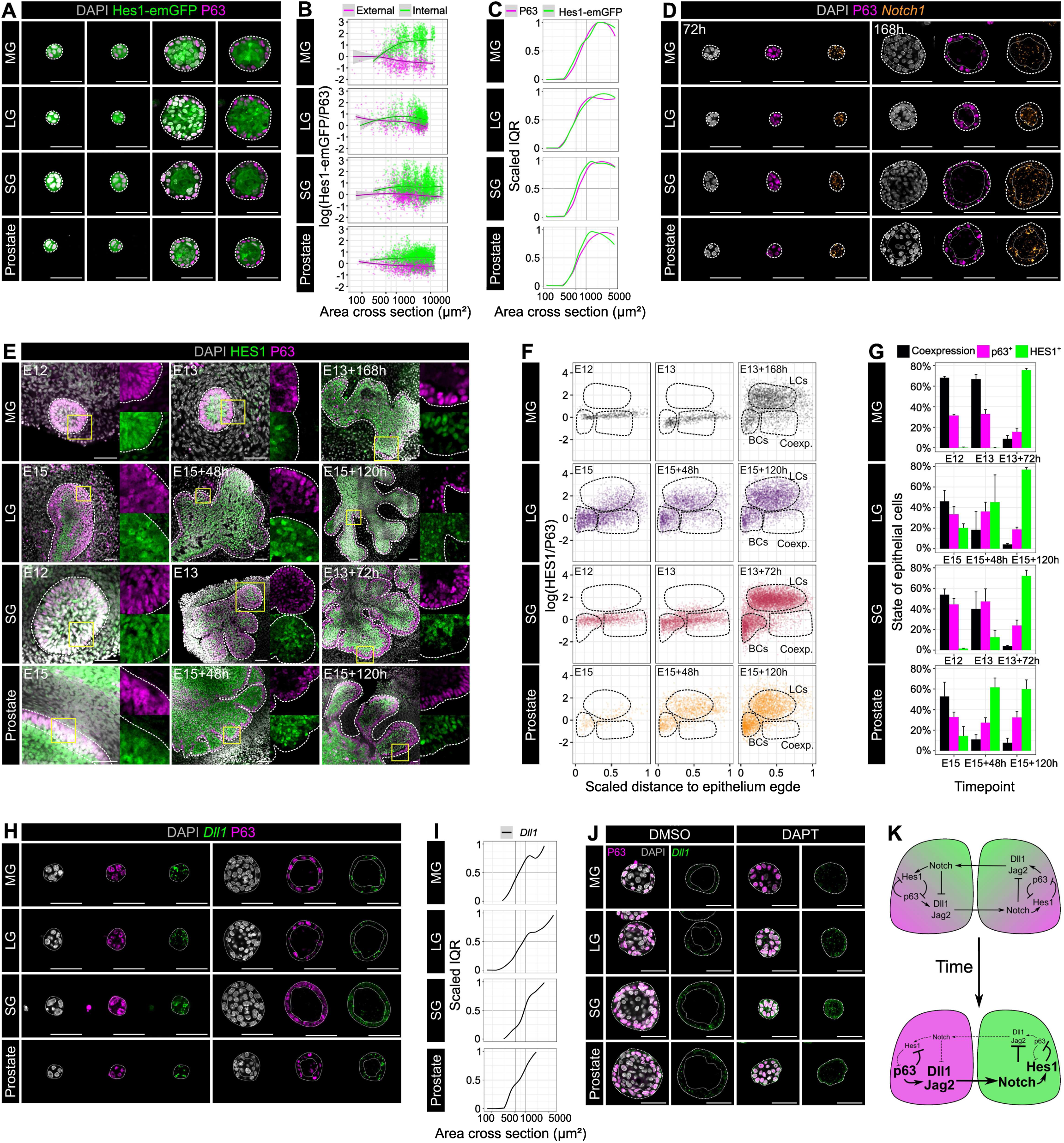
Notch breaks symmetry following a radial geometrical pattern. **A:** Confocal imaging of immunostaining for P63 in Hes1-emGFP organoids at different stages of growth. Scale bar: 50 µm. **B:** Hes1-emGFP and P63 log ratio in cells from organoids of different size (n=3 independent cell sorting experiments). Cells were automatically classified as “Internal” or “External” based on the distance of the nucleus to the external boundary of the organoid. See Methods section for details of image analysis. **C:** Interquartile range (Q3-Q1) of the P63 and Hes1-emGFP intensities measured in each cell of a center optical cross section of organoids of different size (n=3 independent cell sorting experiments). See Methods section. **D:** Confocal imaging of smFISH for *Notch1* and immunostaining for P63 in WT organoids after 48 or 168 h in culture. Scale bar: 50 µm. **E:** Confocal imaging of immunostaining for P63 and HES1 in WT explants at different stages of growth. The dotted line delineates the epithelium limit. Scale bar: 50 µm. **F:** HES1 and P63 log ratio in cells from explants cells at different stage of growth in relation with their normalized distance to the epithelium limit (n=3 biological replicates). **G:** Percentage of HES1^+^, p63^+^ and co-expressing cells at different stage of growth of embryonic explants based on gates displayed in F. Data are displayed as Mean + sd (n=3 biological replicates).**H:** Confocal imaging of smFISH for *Dll1* and immunostaining for P63 in WT organoids at different stages of growth. Scale bar: 50 µm. **I:** Interquartile range (Q3-Q1) of the *Dll1* signal measured in each cell of a center optical cross section of organoids of different size (n=3 independent cell sorting experiments). See Methods section. **J:** Confocal imaging of smFISH for *Dll1* and immunostaining for P63 in WT organoids treated with DMSO or 50µM DAPT for 7 days. Scale bar: 50 µm. **K:** Scheme of gene regulatory network regulating lateral inhibition in bi-layered epithelia.

We subsequently quantified the levels of Hes1-emGFP and p63 in each cell from organoids of different size and calculated the log-ratio of Hes1-emGFP and p63 for each nucleus (Supplementary figure 2C, see Methods). In small organoids (non-segregated fate), we observed a balanced expression of *Hes1* and *Trp63* resulting in a log-ratio around zero. In larger organoids, instead, two cell populations became apparent. The internal position of cells correlated with higher *Hes1* levels while p63 expression was found elevated in external cells (Figure 4B). The segregation of *Hes1* and p63 occurred concomitantly, in organoids within a size range of 500-750 µm², corresponding to 13-21 cells (Figure 4C).

To identify the molecular events that lead to asymmetrical Notch activation, we first characterized the expression of the *Notch1* receptor during organoid growth. As previously observed *in vivo* in the early embryonic MG (Lilja et al, 2018), *Notch1* was homogeneously expressed in all cells from the four tissues as early as 72 hours after culture (Figure 4D), at a stage when epithelial cells still co-express *Trp63* and *Hes1.* To identify cells expressing Notch1 at later stages of development, we derived organoids from *Notch1-Cre^ERT2^:mTmG* mice and treated them with a 24 hour pulse of 4-OHT at day 6 of culture, to label with mGFP the Notch1-expressing cells. Although most mGFP-labeled cells were luminal, we observed, in all four tissues, a minority of K5^+^/mGFP^+^ cells (Supplementary figure 4C), indicating that *Notch1* remains expressed in some BCs after lineage commitment, as confirmed by single molecule fluorescent in situ hybridization (smFISH) in organoids (Figure 4D).

The remaining expression of *Notch1* in BCs prompted us to investigate if Notch pathway activation was the essential trigger of symmetry breaking. This would be consistent with observations in other contexts and model organisms, indicating that Notch receptor expression does not necessarily reflect Notch activation (Lloyd-Lewis et al., 2019; Mukherjee et al., 2005).

To confirm *in vivo* our results in organoids, we investigated the dynamics of expression of *Hes1* in Hes1-emGFP embryonic tissues and by immunostaining against the HES1 protein (Figure 4E, Supplementary figure 4D). We quantified p63 and HES1 levels in embryonic tissues at different developmental stages using a custom image analysis pipeline (Supplementary figure 4E) and found that, in all four tissues, p63 and HES1 were homogeneously co-expressed in embryonic stem cells before lineage specification but were segregated to only one of the two compartments upon commitment to a differentiation program, with p63 in BCs and HES1 in LCs (Figure 4E-G).

We also validated the pattern of *Notch1* expression in embryonic tissues. We labeled Notch1-expressing cells by supplementing the medium of *Notch1-Cre^ERT2^:mTmG* embryonic explants with 4-OHT for 24 hours before fixing the tissues and immunostaining for p63. In this experimental setting, again, we observed a small number of p63^+^/mGFP^+^ cells, thus co-expressing p63 and *Notch1*. This indicates that *Notch1* expression in embryonic explants does not automatically correspond to Notch signal activation, akin to our observations in organoids (Supplementary figure 4F). Together, these findings demonstrate that Notch activity is spatially restricted to internally localized cells during both organoids growth and embryonic tissues development.

### Stem cell commitment follows a lateral inhibition process

To further explore the kinetics of Notch pathway activation, we then investigated the dynamics of expression and localization of the Notch ligands. Mammals encode five Notch ligands belonging to two families, Delta (*Dll1, Dll3* and *Dll4*) and Jagged (*Jag1*, *Jag2*). We explored scRNAseq datasets of each of the four tissues and found that *Dll1*, *Jag1* and *Jag2* were the most highly expressed ligands in all four epithelia (Supplementary figure 4G).

We then assessed the expression of the selected ligands during organoid growth. We observed homogeneous *Jag1* expression, even in organoids with already committed BCs and LCs (Supplementary figure 4H). We concluded that symmetry breaking of Notch activity was unlikely to depend on *Jag1*. On the other hand, *Dll1* and *Jag2* showed a different expression pattern: initially homogeneous in all cells, alongside *Notch1*, *Dll1* and *Jag2* were promptly restricted to external cells as organoids grew (Figure 4H, Supplementary figure 4I). We quantified *Dll1* and *Jag2* expression for each cell within organoids (Supplementary figure 4-J, see Methods) of different size and observed that the critical threshold for segregation of ligand expression corresponded to a size range between 500-750 µm² (∼13-21 cells), consistent with the time of segregation of p63 and Hes1 expression (Figure 4I, Supplementary figure 4K).

We then assessed the *in vivo* expression of *Dll1* and *Jag2* by data mining published scRNAseq data of embryonic MG, SG and prostate (Carabaña et al., 2024; Lee et al., 2021; Wang et al., 2021), whereas the E16 LG (Farmer et al., 2017) dataset was composed of too few epithelial cells to perform such analysis. We observed a clear enrichment of *Dll1* and *Jag2* in BC-like cells right at the time of fate specification, while *Jag1* and *Notch1* remained expressed in all epithelial cells (Supplementary figure 4L). These data indicated a clear correlation between restriction of expression of the Notch ligands *Dll1* and *Jag2* to external cells and onset of symmetry breaking, both in organoids and in embryonic tissues.

Notch is known to regulate pattern formation by a process called lateral inhibition (Heitzler and Simpson, 1991). Lateral inhibition occurs when small variations of Notch signaling are amplified by receptors-ligands feedback loops. To understand how the differential pattern of Notch activity and ligands expression is established in our tissues, we then investigated the effect of Notch inhibition on the expression of *Dll1* and *Jag2*. After 7 days in culture in the presence of DAPT, organoids failed to segregate both *Dll1* and *Jag2* expression into external cells, demonstrating the existence of a negative feedback loop between Notch activation and *Dll1* and *Jag2* expression (Figure 4J, Supplementary figure 4M). Once symmetry of Notch activity brakes through ligand-receptor trans-activation, the position-dependent cell heterogeneity is further amplified by Notch-induced repression of *Dll1/Jag2* expression and previously described *p63* downregulation (Nguyen et al., 2006; Tadeu and Horsley, 2013). Ligand-receptor cis-interaction prevent Notch activation in external cells previously called cis-inhibition (Álamo et al., 2011). p63 in turn reinforces the patterning of ligands through activation of the expression of *Dll1* and *Jag2* and repression of *Hes1* (Melino et al., 2015; Yalcin-Ozuysal et al., 2010). Based on these observations, we propose that Notch activation breaks symmetry following a classical lateral inhibition model (Figure 4K).

### YAP activity is spatially patterned in organoids and embryonic development

A homogeneous group of cells breaks symmetry through a lateral inhibition mechanism if ligands expression is variable enough. In this case, theory predicts that symmetry could break in organoids as small as two cells. Contradictory to this theoretical prediction, we found experimentally that organoids of the four tissues remain homogeneous until they reach the critical size of 500-750 µm² (13-21 cells). We therefore postulate that an active mechanism must exist to maintain homogeneous Notch activity at early times (i.e. in small organoids).

Given the observed correlation with organoid size and tissue architecture, we interrogated the expression and activity of YAP, a transcription factor involved in mechano-transduction downstream of the Hippo signaling pathway. We therefore quantified the nuclear to cytoplasmic ratio of YAP within each cell composing an organoid and observed that YAP activity displayed a spatial pattern dependent on organoid size and cell position. In small organoids, YAP was homogeneously localized in the nucleus of all cells (Figure 5A, B). However, as soon as cells became internally localized, we observed nuclear YAP only in external cells (Figure 5A, B). To confirm that nuclear YAP localization coincided with YAP activity, we evaluated the expression of *Ctgf*, a known direct target of YAP signaling and observed exclusive localization of *Ctg*f transcript in external cells displaying nuclear YAP (Supplementary figure 5A). We then assessed YAP localization relative to p63 and HES1, by co-staining for the three proteins at different stages of organoid growth (Figure 5C). For each cell imaged, we extracted the intensity values of Hes1-emGFP and p63 and the YAP nuclear to cytoplasmic ratio, alongside the organoid area and the cell position within the organoid, and we used principal components analysis (PCA) to reduce the dimensionality of our data (Figure 5D). Cells within organoids smaller than 500 µm² clustered together, while two other distinct clusters emerged, composed of either internal or external cells. Internal cells displayed high HES1 levels and low YAP activity, characteristic of LCs, while external cells displayed strong p63 signal and nuclear YAP, reminiscent of BCs identity (Figure 5D, supplementary figure 5B).

**Figure 5.**
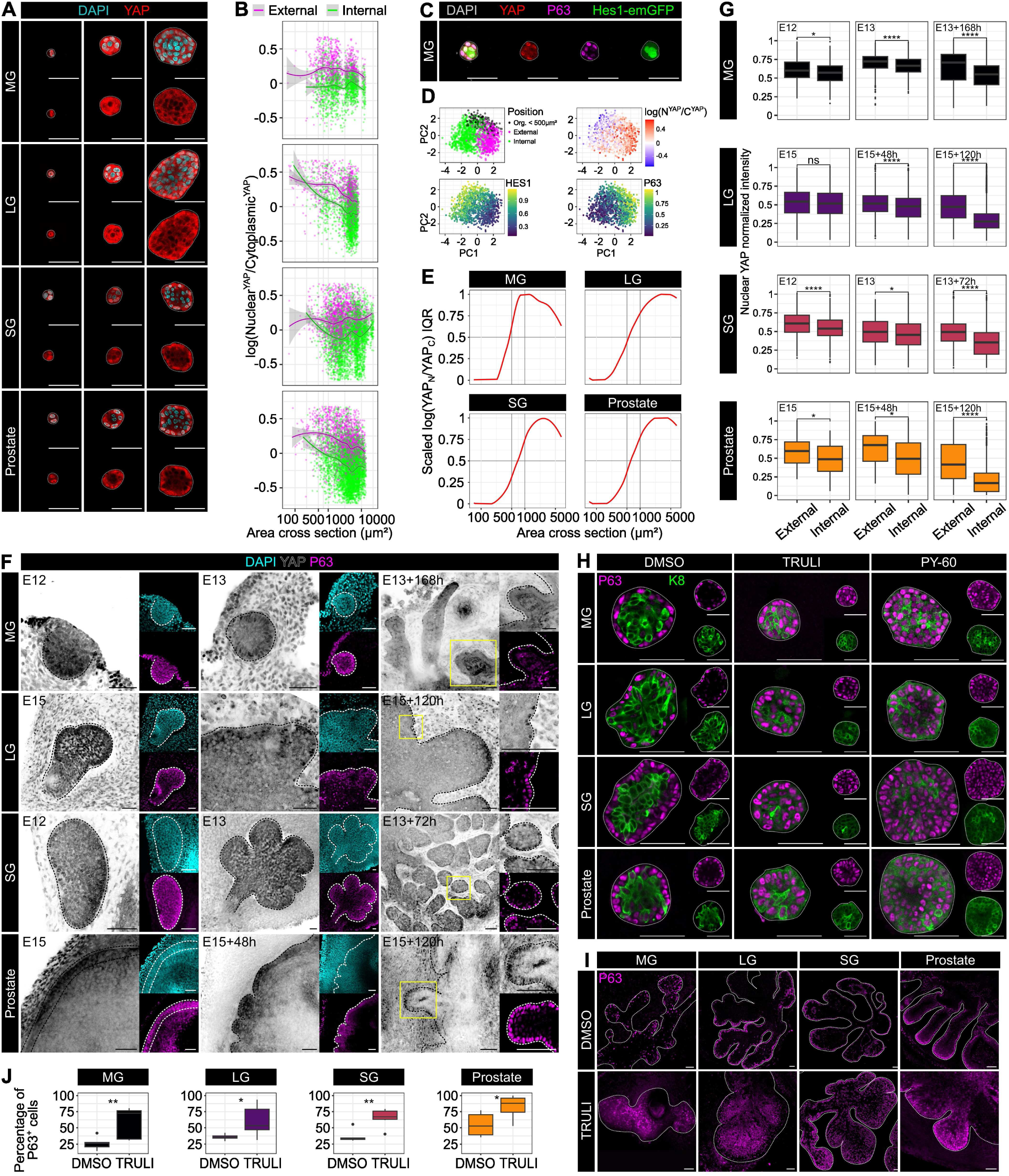
YAP activity is spatially patterned during organoid growth and embryonic development. **A:** Confocal imaging of immunostaining for YAP in WT organoids at different stages of growth. Scale bar: 50 µm. **B:** YAP nuclear to cytoplasmic log ratio in cells from organoids of different size (n=3 independent cell sorting experiments). Cells were automatically classified as “Internal” or “External” based on the distance of the nucleus to the external boundary of the organoid. See Methods section for details of image analysis. **C:** Confocal imaging of immunostaining for P63 and YAP in Hes1-emGFP organoids. Scale bar: 50 µm. **D:** Principal component analysis (PCA) of cells stained for YAP and P63 in Hes1-emGFP MG derived organoids of different size (n=3 independent cell sorting experiments). See Methods section. **E:** Interquartile range (Q3-Q1) of the YAP intensity measured in each cell of a center optical cross section of organoids of different size (n=3 independent cell sorting experiments). See Methods section. **F:** Confocal imaging of immunostaining for P63 and YAP in WT explants at different stages of growth. Scale bar: 50 µm. **G:** Quantification of normalized nuclear YAP intensity in internal or external cells of WT explants at different stages of growth (n=3 biological replicates). Non-parametric Wilcoxon test. **H:** Confocal imaging of immunostaining for p63 and K8 in WT organoids treated with DMSO, TRULI or PY-60 for 7 days. Scale bar: 50 µm. **I:** Confocal imaging of immunostaining for P63 in WT explants treated with DMSO or TRULI for the whole duration of the culture. Scale bar: 50 µm. **J:** Percentage of P63+ cells in explants treated with 20 µM TRULI for the whole duration of the culture (n=3 biological replicates). Data displayed as boxplot. Non-parametric Wilcoxon test.

We then assessed the timing of symmetry breaking of YAP activity by measuring the interquartile range of YAP nuclear to cytoplasmic log ratio. Of interest, we found that symmetry breaking of YAP activity also occurred around 500-750 µm² (Figure 5E), coinciding with lineage segregation of the early fate markers p63 and Hes1. To establish the relevance of our findings in embryonic development, we stained embryonic tissues for YAP at different developmental times and observed a progressive segregation of YAP activity to external cells, corroborating our findings in organoids (Figure 5F, G). These results indicate that YAP activity is spatially patterned during organoid growth and embryonic tissue development. Initially homogeneously active in embryonic epithelial stem cells, YAP remains only in the nucleus of external cells at the onset of fate specification.

### YAP regulates the binary fate decisions of bi-layered epithelial stem cells

To investigate the role of YAP in binary cell fate decisions, we then grew organoids in the presence of drugs affecting YAP signaling. We first tested the effect of different doses of Verteporfin, a potent inhibitor of the YAP-TEAD interaction (Wang et al., 2015) and of two other YAP inhibitors, MGH-CP1 (Li et al., 2020) and TED-347 (Bum-Erdene et al., 2019). All these inhibitors led to rapid cell death impairing organoid growth (Supplementary figure 5C), indicative of the critical role of YAP in our system but precluding further analysis of its specific role in cell fate asymmetry. We therefore used a complementary approach and forced YAP activation and nuclear localization by treating organoids with TRULI, a *Lats1/2* kinase inhibitor, or with PY-60, a molecule recently described to increase nuclear accumulation of YAP by inhibiting its cytoplasmic retention (Kastan et al., 2021; Shalhout et al., 2021). We first established that YAP nuclear localization and activity was indeed promoted by treatment with these compounds, using *Ctgf* RNA expression as a readout for YAP activity (Supplementary figure 5D). We then submitted the organoids to a continuous treatment with TRULI or PY-60. After 7 days, organoids cultured in the presence of YAP activators were composed of a homogeneous population of cells co-expressing P63 or K5 and low levels of K8 (Figure 5H, Supplementary figure 5E), demonstrating that homogenous YAP activity prevents fate specification. We also examined the effect of TRULI on YAP localization in embryonic explants (Supplementary figure 5F) and found that YAP activation in embryonic explants led to a dramatic increase in the number of p63^+^ cells, accompanied by impairment of branching morphogenesis in all four tissues examined (Figure 5I-J). Together, these experiments demonstrate that YAP activity must be spatially patterned for proper fate specification to occur.

### Active YAP prevents symmetry breaking and maintains a stem-like cell state

To further characterize the effect of YAP activation on cell fate and establish its relationship to Notch activity and p63 expression, we then performed immunostaining for p63 and HES1 in large organoids cultured in the presence of TRULI or PY-60 (Figure 6A, B). As anticipated, p63 and HES1 were mutually exclusive in control conditions (DMSO) and the Notch inhibitor DAPT completely prevented the formation of LC-like internal cells. However, TRULI or PY-60 treatment led to a striking increase in the number of cells co-expressing p63 and HES1, even in large organoids, where the two proteins are normally segregated, and cells are already committed to a specific lineage (Figure 6A, B). The ectopic co-expression of p63 and HES1 in large organoids is reminiscent of the homogeneous distribution pattern of these fate determinants before symmetry breaking, at early stages of organoid growth (Supplementary figure 6A) or during embryonic tissue development (Figure 4E). Based on these unexpected findings, we therefore postulate that YAP activation prevents symmetry breaking and promotes the maintenance of an embryonic stem-like state. To confirm this hypothesis in embryonic development, we treated embryonic explants with TRULI and observed an increase in the number of cells co-expressing p63 and HES1 (Figure 6C, D, Supplementary figure 6C).

**Figure 6.**
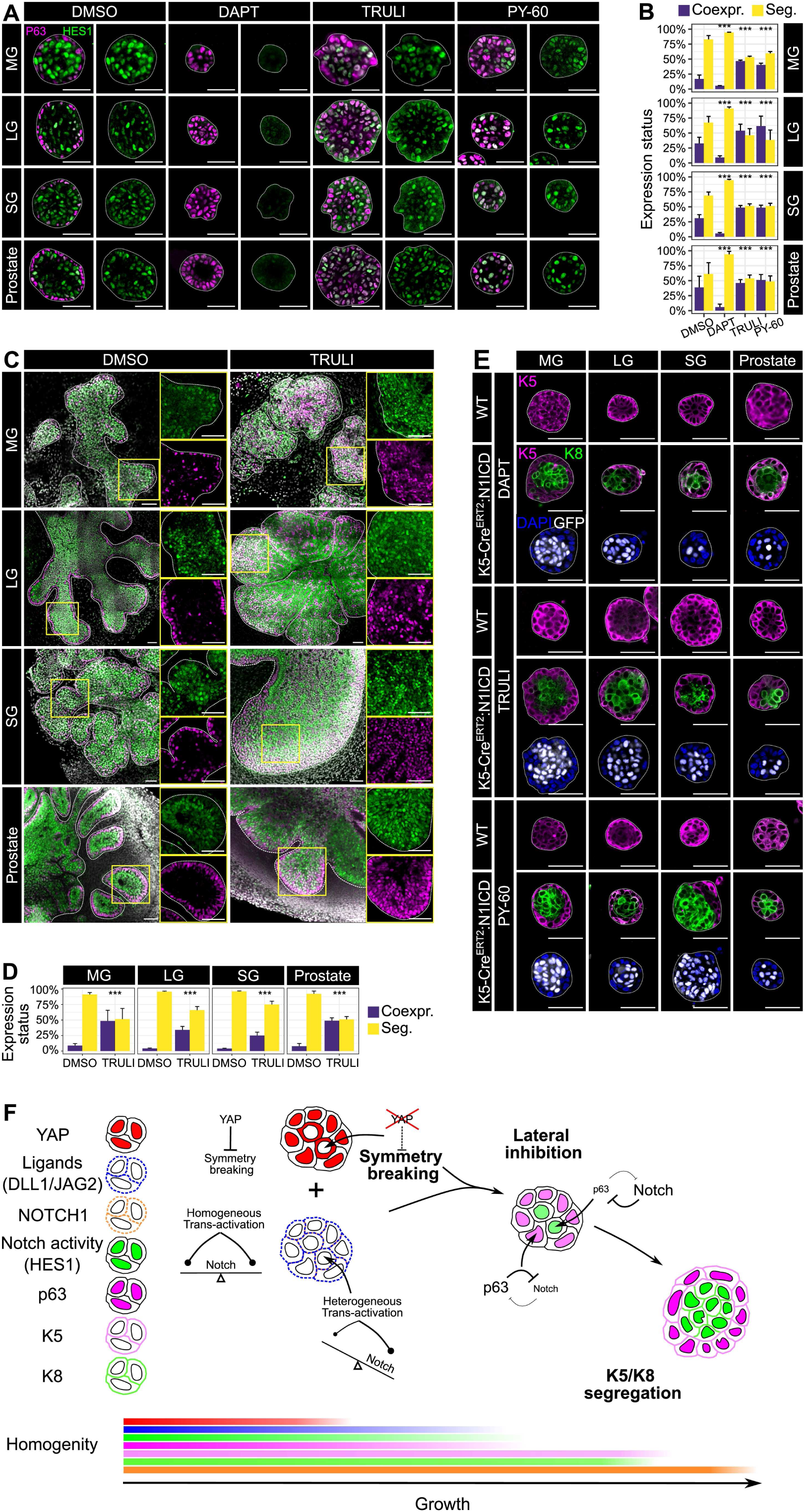
Active YAP prevents symmetry breaking and maintains a stem-like cell state. **A:** Confocal imaging of immunostaining for p63 and HES1 in WT organoids treated for 7 days with DMSO, 50 µM DAPT, 20 µM TRULI or 10 µM PY-60. Scale bar: 50 µm. **B:** Quantification of the percentage of cells co-expressing P63 and HES1. Data are displayed as Mean + sd (n=3 independent cell sorting experiments). Chi square test. **C:** Confocal imaging of immunostaining for p63 and HES1 in WT explants treated with DMSO or 20 µM TRULI for the whole duration of the culture. Scale bar: 50 µm. **D:** Quantification of the percentage of cells co-expressing P63 and HES1 in explants treated with DMSO or TRULI for the whole duration of the culture (n=3 biological replicates). Data are displayed as Mean + sd. **E:** Confocal imaging of immunostaining for K5, K8 and nGFP in WT or *K5-Cre^ERT2^:N1ICD-nGFP* organoids treated for 7 days with DMSO, 50 µM DAPT, 20 µM TRULI or 10 µM PY-60. Scale bar: 50 µm. **F:** Model of binary fate decision in single cell derived organoid.

Our initial quantification of the extent of cellular heterogeneity within organoids of different size allowed us to observe a clear difference between the timing of fate commitment and the segregation of the lineage-specific cytokeratins K5 and K8 but did not have the temporal resolution to determine whether p63, YAP, HES1 or Notch ligands break the symmetry first (Supplementary figure 6B). Therefore, to determine the very first event of symmetry breaking we probed the epistatic relationship between YAP and Notch. We treated organoids derived *from K5-Cre^ERT2^:N1ICD* mice with the YAP activators TRULI or PY60 and assessed if mosaic ectopic Notch activation in uncommitted cells could rescue the phenotype induced by YAP activation. We first confirmed that expression of the cleaved form of the Notch1 receptor was sufficient to generate LCs even in the presence of DAPT (Figure 6E, Supplementary figure 6D). Similarly, *Notch1* gain-of-function could also rescue the defect in LCs production caused by ectopic YAP activation (Figure 6E, Supplementary figure 6D). To corroborate these results, we performed immunostaining for YAP in organoids composed only of N1ICD-expressing mutant cells (nGFP+) which occurs when mutation is induced at the 1 cell stage, and we detected externally localized cells with nuclear YAP and K8 expression (Supplementary figure 6E).

Together, these data indicate that YAP activity promotes a stem-like state and the absence of spatial patterning of YAP activity impairs symmetry breaking. Furthermore, the epistatic observation that ectopic Notch activation can overcome the restriction in symmetry breaking caused by homogeneous YAP activity, indicates that YAP acts upstream of Notch, delaying lateral inhibition until a cell becomes internally localized (Figure 6F).

### YAP, p63 and HES1 are co-expressed during regeneration

The organoid model we have employed here relies on reactivation of multipotency of adult committed basal cells, mimicking to some extent tissue regeneration. Therefore, we investigated the conservation of the Notch-p63 signaling axis *in vivo* during adult tissue regeneration. We first probed the mechanisms underlying reactivation of multipotency of BCs following LCs ablation in the MG and the prostate, as described by Centonze and colleagues (Centonze et al., 2020). In this study, the authors elegantly combined BC lineage tracing with genetic depletion of LCs in adult *K5-Cre^ERT2^/R26-tdTomato/K8rtTA/TetO-DTA* mice and observed reactivation of BCs multipotency (Figure 7A) (Centonze et al., 2020). scRNAseq of MG cells after LCs ablation indicated that multipotent BCs enter a hybrid state where they co-express basal and luminal markers, including *Krt8*, *Krt18*, *Krt5*, *Krt14* and *Trp63* (Figure 7B, Supplementary figure 7A-D) (Centonze et al., 2020). Co-expression of opposite lineage markers is reminiscent of the uncommitted cell state we describe here during organoid growth and *in vivo* in embryonic development. We thus evaluated the expression level of *Trp63* and *Hes1* in 3 different MG populations: adult LCs, BCs and hybrid cells. We observed that the hybrid MG cluster is enriched in cells co-expressing Trp63 and *Hes1*, while BCs and LCs express exclusively one or the other lineage marker (Figure 7C). These hybrid cells expressed also *Notch1*, *Hes1* and the Notch ligands *Dll1, Jag1* and *Jag2*, at intermediate levels (Supplementary figure 7E). To corroborate these in silico observations, we performed immunostaining for p63 and HES1 on MG and prostate tissues upon LC ablation and could find cells co-expressing the two opposing fate determinants 1 week after LCs ablation, while p63 and HES1 were clearly segregated in control mice (Figure 7D). Furthermore, we assessed YAP activity following LCs ablation. While nuclear YAP was predominantly found in BCs in normal MG and prostate tissue, upon LCs ablation, we observed an overall increase in nuclear YAP, that could also be detected in LCs (Figure 7E). Sequencing data confirmed this observation, showing elevated levels of *Yap1* and its target *Ctgf* in hybrid cells (Supplementary figure 7F).

**Figure 7.**
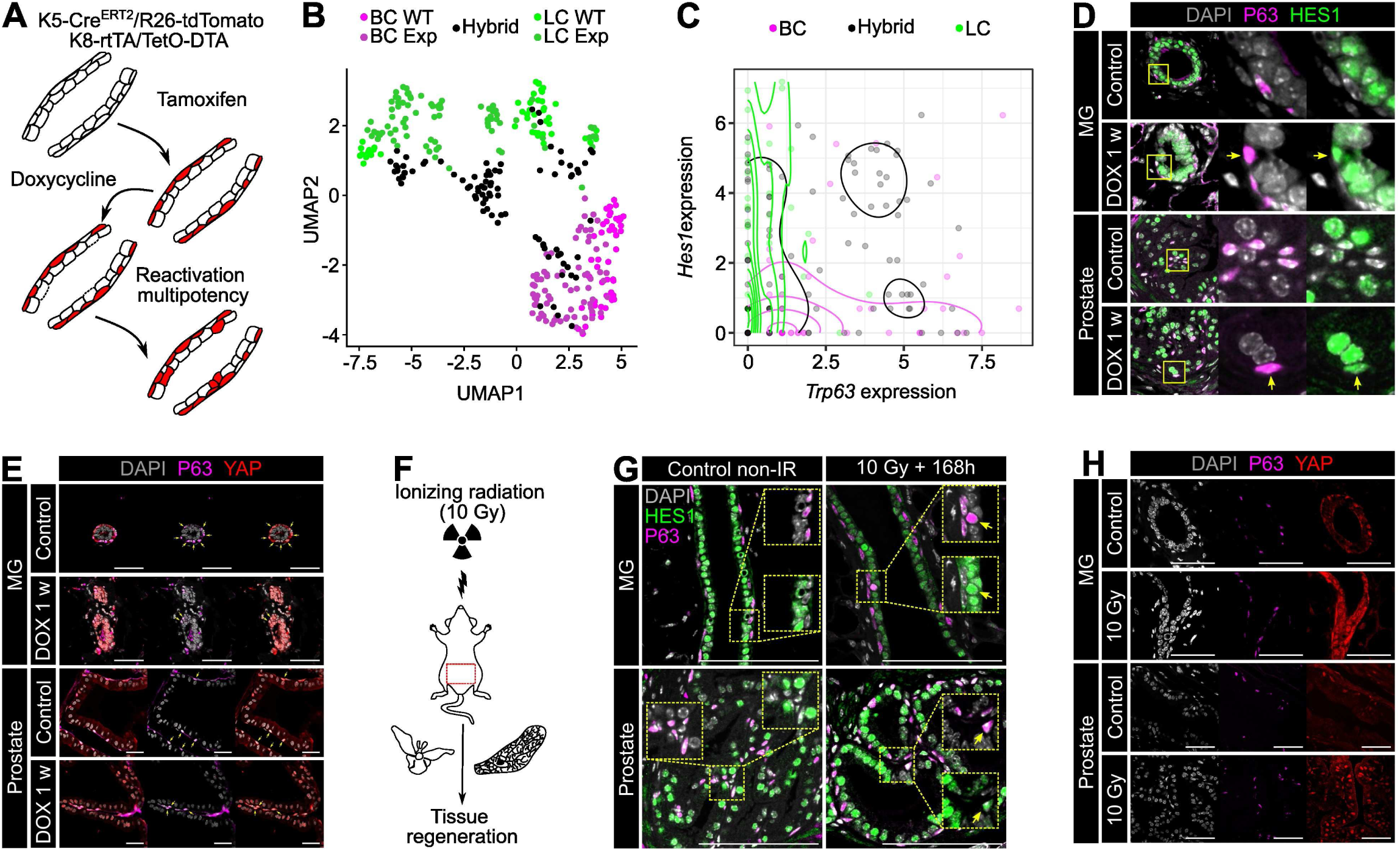
YAP, p63 and HES1 are co-expressed during regeneration. **A:** Scheme of the ablation induced reactivation of multipotency in BCs used in Centonze et al. **B:** Dimensionality reduction using UMAP of scRNA-seq data of MG epithelial tissues cells sequenced 1-week post ablation and after labelling of BCs. Cell population identified during sorting are represented color-coded (Centonze et al., 2020). **C:** Co-expression of Trp63 and Hes1 in the WT LC, BC or Hybrid (Tom^+^) cell populations. Density and cell type cells are color-coded. **D:** Confocal imaging of immunostaining for P63 and HES1 in tissue before ablation (Control) or 1 week after ablation (DOX 1 w). **E:** Confocal imaging of immunostaining for YAP and P63 in non-ablated tissue (Control) or 1 week after ablation (DOX 1 w). **F:** Scheme of the irradiation induced BCs reactivation of multipotency approach. **G:** Confocal imaging of immunostaining for P63 and HES1 in adult tissue before irradiation (Control non-IR) or 1 week after (10 Gy +168h). **H**: Confocal imaging of immunostaining for p63 and YAP in adult tissue before irradiation (Control non-IR) or 1 week after (10 Gy +168h).

To further validate our findings in another regenerative context, we employed ionizing radiation, a widely used technique in various tissues (Serrano Martinez et al., 2021). We investigated the regeneration dynamics of the mammary and prostatic glands in adult mice after a dose of 10 Gy of ionizing radiation on the lower abdomen (Figure 7F). We used the gut of the irradiated animals as a positive control to recapitulate the previously described dynamic of cell death, proliferation, YAP and HES1 during intestinal regeneration (Yui et al., 2018) (Supplementary figure 7G). We then investigated the dynamics of cell death and proliferation in MG and prostate and observed a peak of cell death between 1 and 3 days after irradiation and an increase of proliferation from 3 days to 7 days post-irradiation (Supplementary figure 7H, I). We then immunostained the irradiated tissues for HES1 and p63 seven days post irradiation, alongside non-irradiated controls. While we confirmed the clear segregation of p63 and HES1 in non-irradiated tissues, we identified cells that were co-expressing the two fate markers 7 days post-injury (Figure 7G). In addition, we observed an expanded population of cells presenting nuclear YAP in irradiated tissues, mirroring the results observed upon LCs ablation (Figure 7H).

These data suggest that the reactivation of multipotency during adult tissue regeneration is reflected by a hybrid cell state exhibiting co-expression of basal and luminal markers. Our results suggest that this plastic metastable state is maintained by the simultaneous activation of p63 and Notch signaling, orchestrated by YAP activity. Overall, this state evokes and re-uses fetal developmental programs of cell fate specification characterizing embryonic multipotent stem cells.

## Discussion

In agreement with previous work reporting that YAP activity promotes stemness and self-renewal in other epithelia (Panciera et al., 2016; Schlegelmilch et al., 2011), we found that YAP is initially homogeneously active in uncommitted stem cells of the MG, SG, LG and prostate. These embryonic-like stem cells brake symmetry when organoids reach a critical size that corresponds to 13 cells. The cell that first excludes YAP from the nucleus is invariably located at the center of the organoid. Interestingly, 12 is the number of spheres of the same size that can be arranged in space while maintaining contact with a common central sphere, a property also known as the “kissing number” (Pfender and Ziegler, 2004). The global structure of our spherical organoids, composed of 13 cells, accurately reflects the conformation of a ball of spheres.

Molecularly, we found that symmetry breaking of bi-layered epithelial cells follows a classical lateral inhibition model. While such model would predict that symmetry could break already at the 2-cell stage, we observed experimentally that Notch and p63 symmetry breaking occurs when organoids reach the same number of 13 cells. Similarly to what has been reported in intestinal organoids (Serra et al., 2019), ectopic YAP activation in both organoids and embryonic tissues blocks symmetry breaking of Notch and p63 and, consequently, stem cell commitment. These observations prompt us to propose a model in which YAP acts as a capacitator, integrating the information of the cell position within organoids and delaying symmetry breaking to allow commitment only when some cells become internally localized. The first internalized cell then excludes YAP from the nucleus and activates Notch signaling, possibly via its extensive cell-cell contact area with surrounding cells expressing the Notch ligands (Shaya et al., 2017). This event is the initial asymmetric step enabling lateral inhibition.

The molecular crosstalk between Hippo/YAP and Notch pathways has been proposed to regulate a wide range of developmental processes, with different context-dependent modalities of interactions (Totaro et al., 2018). Indeed, Notch can cooperate (in trophectoderm specification) (Rayon et al., 2014), act upstream (in cortical development) (Liu et al., 2018) or downstream (during epidermis differentiation) (Totaro et al., 2017) of the Hippo/YAP pathway. In the four tissues examined here, our epistatic experiments of concomitant activation of Notch and YAP (Figure 6-E, Supplementary figure 6-E) support a model where Notch acts downstream of YAP. Interestingly, YAP has been previously reported to interact with both p63 and HES1 and to regulate their respective transcriptional targets (Mamidi et al., 2018; Zhao et al., 2014). Based on our findings, we can propose a mechanism by which YAP promotes stemness by balancing the activity of the mutually antagonistic p63 and Notch pathways.

In addition, our results suggest that the Hippo/YAP pathway depends on the cell position within growing organoids or tissues. It is well established that the YAP pathway can be regulated by biomechanical signals such as cell density, substrate stiffness or cell polarity (Totaro et al., 2018). In our organoids, we observed the appearance of a geometric radial heterogeneity concomitantly with the segregation of active YAP to external cells (Supplementary figure 6-J), suggesting that positional cues may influence YAP patterning in the tissues studied here; however, further investigations are needed to precisely define which cues regulate YAP symmetry breaking.

The minimum number of epithelial cells required for symmetry breaking differs between organoid and embryonic development. Indeed, *in vivo*, embryonic buds composed of more than 13 epithelial cells still co-express YAP, P63 and HES1. This difference may be explained by the different environmental cues to which epithelial cells are exposed to in the embryo and in organoids, which could affect YAP activity. However, the overall model of symmetry breaking regulation by positional cues and tissue geometry remains valid since we found that YAP is initially excluded from the nucleus of the inner cells in both organoids and mouse embryos.

In all tissues examined, YAP remained active in some adult BCs, which are characterized by a higher degree of plasticity compared to adult LCs, as exemplified by their greater ability to revert to a multipotent stem-like cell state when homeostasis is disrupted. The unipotent behavior of homeostatic adult BCs, despite YAP activation, has been shown to be maintained by cell-cell contact between BCs and LCs (Centonze et al., 2020).

The cell state promoted by YAP activity, characterized by the co-expression of opposing lineage markers (p63 and HES1), is reminiscent of the hybrid cell state observed after cell-cell contact disruption in MG and prostate regeneration. This cell state evokes embryonic/fetal transcriptional programs, that can be triggered by YAP activation during regeneration in other tissues (Gregorieff et al., 2015; Pardo-Saganta et al., 2015; Pikkupeura et al., 2023; Serra et al., 2019; Yui et al., 2018). Our data, consistent with the previous studies cited above, thus demonstrate the broadly conserved role of YAP in establishment and maintenance of epithelial tissue integrity and function.

In conclusion, in agreement with very recent studies (Alasaadi et al., 2024; Shroff et al., 2024), we show that tissue architecture and geometry can regulate molecular pathways to drive the establishment of fate patterns, and that YAP activity represents a central player in coordinating the temporal and spatial dimensions during tissue morphogenesis and regeneration.

Finally, the remarkable conservation of these critical mechanisms in four different bi-layered epithelia originating from separate germ layers stresses their importance for tissue development and regeneration. Just as the phylotypic stages during animal embryogenesis, the fate specification mechanisms that we reveal here seem to follow an hourglass model, where YAP, Notch and p63 represent the core conserved players instructing cell fate in tissues of different origin that will later diverge to acquire tissue-specific features.

## Methods

### Statistics and Reproducibility

Experiments were performed in biological and technical replicates as indicated. A biological replicate corresponds to pups from different litters for embryonic experiment. Siblings included in a same experiment are considered as technical replicates. Organoids originating from independent cell sorting experiments were considered as biological replicates. Each independent cell sorting experiments contains cells from 2 to 4 animals of the same genotype pooled after dissection. Experiments with at least 3 biological replicates were used to calculate the statistical value of each analysis. All graphs are mean + SD. Statistical analysis was performed using the Wilcoxon test or Chi squared test.

### Ethics Statement and mouse models

All studies and procedures involving animals were carried out in strict accordance with the recommendations of the European Community (2010/63/UE) for the protection of vertebrate species used for experimental and other scientific purposes. Approval was granted by the Ethics Committee of the Institut Curie CEEA-IC #2021-029 and the French Ministry of Research (reference #34364-202112151422480). We adhere to the internationally established principles of replacement, reduction and refinement according to the Guide for the Care and Use of Laboratory Animals (NRC 2011). Animal husbandry, supply, maintenance and care in the animal facility of the Institut Curie (facility license #C75-05-18) before and during experiments fully met the needs and welfare of the animals. All animals were housed in individually ventilated cages under a 12:12h light/dark cycle, with water and food available ad libitum. All mice were culled by cervical dislocation. Mice were genotyped by PCR on genomic DNA extracted from an earpiece for adult mice or tail tip for embryos. Plug detection at mid-day was considered 0.5 days post-coitus (E0.5). Mouse breeding and husbandry was managed using the mouse colony organization software MiceManager: https://infenx.com/mouse-colony-management-software/. See Key Resources Table 3 for a complete list of mice used in this work.

Organoids and explants were derived from the WT, *mTmG* double fluorescent reporter line (Muzumdar et al., 2007). The K5-Cre^ERT2^ mouse (Van Keymeulen et al., 2011) was crossed with *R26:mTmG* or *N1ICD-nGFP* animals (Murtaugh et al., 2003) to study the effect of Notch activation on fate commitment in organoids. We crossed the *R26-Cre^ERT2^* line (Ventura et al., 2007) with *mTmG* or *N1ICD-nGFP* for embryonic experiments. We used the *Hes1-emGFP* reporter mouse animals to monitor Notch activity (Fre et al., 2011) and the *Notch1-Cre^ERT2^* animals to label Notch1 expressing cells (Fre et al., 2011). All mice used were from a mixed genetic background. Reporters and mutant alleles were recombined by a single intraperitoneal injection of tamoxifen free base prepared in sunflower oil containing 10% ethanol (0.1 mg per g of mouse body weight), unless indicated otherwise in figure legends.

### Tissue irradiation

10-12 weeks old C57BL6/J mice were treated with conventional electrons using a wide field of12,7 x 5,4 cm to allow complete irradiation of prostate for males and mammary gland for females. Anesthesia was carried out with a nose cone using 2.5% isoflurane in the air without adjunction of oxygen. Dosimetry was individually controlled using Gafchromic films positioned on the surface of the mice’s abdomens at the center of the irradiation field.

### Single cell isolation, flow cytometry and sorting

MG, LG, SG and prostate were isolated from 8–12-week-old mice. A typical experiment was generated using two males and two females, with MG and Prostate from two animals pooled, while LG and SG from four animals were pooled and the four tissues were dissociated using the same protocol as previously described (Di-Cicco et al., 2015; Stingl et al., 2006; Taddei et al., 2008). Briefly, single cell dissociation was performed by enzymatic digestion with 600 U ml/collagenase and 200 U/ml for 90 min at 37°C with shaking (160 rpm). Cells were further dissociated in hot trypsin for 1 min, in 5 mg/mL dispase and 0.1 mg/mL DNase I for 3 min at 37°C, then red blood cells were eliminated in 0.63% NH_4_Cl. The cell suspension was filtered through a 40μm cell strainer to obtain a single cell preparation for FACS. Single cell suspensions were incubated for 30 minutes on ice with conjugated primary antibodies diluted in PBS (APC anti-mouse CD31, APC anti-mouse Ter119, APC anti-mouse CD45, APC/Cy7 anti-mouse CD49f, PE/Cy7 anti-mouse EpCAM at 1:100 dilution) (Crowell et al., 2019; Stingl et al., 2006; Yasuhara et al., 2022). Cells were washed with PBS, resuspended in flow buffer containing DAPI and passed through a 40 μm cell filter. FACS analysis was performed on an ARIA flow cytometer (BD). Doublets, dead cells (DAPI+) and Lin+ (CD45+, CD31+, Ter119+) cells were systematically excluded from the analysis. Full details of the antibodies used can be found in Key Resources Table 1. Results were analyzed using FlowJo software (V10.0.7).

To isolate and sort organoid cells, two wells of organoids were harvested in ice-cold PBS and the pellets were then incubated on ice for 15 minutes in cold Cell Recovery Solution to dissolve the Matrigel. Organoids were then centrifuged at 400 × g for 5 minutes at 4°C. Cell dissociation was performed as previously described for tissues.

### *In vitro* organoid cultures

Sorted BCs were embedded in 90% Matrigel and cultured in 24-well plates or 8-well imaging chambers (Ibidi 80827) for whole mount staining. After polymerization, the Matrigel drops were covered with tissue-specific organoid medium containing a base of DMEM/F12 supplemented with 1x Glutamax, 1x HEPES, 1X B27 supplement, recombinant Noggin and R-spondin, stabilized FGF2 and 2% penicillin/streptomycin. MG medium was supplemented with 1x N2 supplement and Neuregulin, SG and LG medium were supplemented with NAC, Wnt3a and EGF and prostate medium was supplemented with dihydrotestosterone (DHT), EGF and A8301 (Crowell et al., 2019; Jardé et al., 2016; Kim et al., 2021). All organoid cultures were supplemented with Y27632 for the first 48 hours of culture. Factors and inhibitors are listed in Key Resources Table 1.

### Immunostaining

Tissues samples were fixed in 4% PFA (2 h at RT) and incubated overnight in 30% sucrose at 4°C before embedding in optimal cutting temperature (OCT) compound. 8 μm thick cryosections were cut using a cryostat (Leica CM1950) and stored at -20°C. For paraffin immunostaining, tissues were fixed overnight at 4°C in 4% PFA, washed 1 h in PBS and incubated in 70% ethanol until embedding. 5 μm thick cryosections were cut using a rotational microtome (Thermo scientific, Microm HM340E) and stored at RT. Section were deparaffinized, rehydrated and boiled in pH6 citrate buffer for 15 minutes to retrieve antigen. Paraffin and frozen sections were blocked (PBS, 5% FBS, 1% BSA) for 1 hour and incubated with primary antibodies diluted in blocking buffer overnight in a humidified chamber. After washing, sections were incubated with a solution of secondary antibodies and DAPI (Merck) diluted in PBS for 1 hour. Finally, the sections were mounted on a slide (Aqua-Polymount, Polysciences).

Whole mount immunostaining was performed on organoids and explants. Organoids were fixed in ice-cold 4% PFA (10 minutes), permeabilized with PBS 1% Triton x100 (1 hour) for organoids and with PBS 5% Triton x100 (24 hours), blocked in PBS, 5% FBS, 1% BSA (1 hour) and incubated with primary antibodies overnight at room temperature (RT) for organoids or two nights at 4°C for explants. After removal of the primary antibody solution, dilutions of secondary antibodies and DAPI in PBS were added to organoids for 6h at RT and overnight at RT for explants. Explants were mounted on a microscope slide using Aqua-Polymount. Organoids were stored in a 1:1 ratio of PBS and glycerol solution. Explants and organoids were imaged by confocal microscopy. A list of all antibodies used here is provided in Key Resources Table 2.

### In situ hybridization

smRNA-FISH was performed using the RNAscope Multiplex Fluorescent Reagent Kit v2 (ACD) according to the manufacturer’s recommendations using commercially available probes. FISH was performed on wholemount organoids in the imaging chamber after fixation (10 min ice-cold PFA 4%) and permeabilization (PBS 1% Triton x100, 1h). Full details of the RNAscope probes used here can be found in Key Resources Table 1.

### Relation between cross section area and number of cells composing organoids

Measure of the roundness of organoids was performed on mask resulting from organoids segmentation, using the formula: 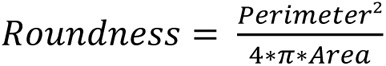

z-stack images of organoids were acquired at different stages of growth and the number of cells composing each organoid was manually counted. The relationship between the area of the central cross section of a sphere and its total volume follows the form:

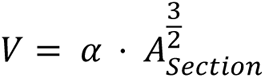

With *V* the volume of the sphere and *A*_*Section*_ the area of the central cross section.

Since the structure formed by organoids is very spherical, we fitted the data obtained with an equation of the same form and obtained the following equation: 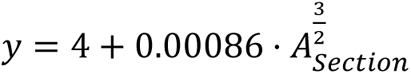

### Image acquisition and analysis

Images were acquired using a LSM780, LSM880 or LSM900 inverted laser scanning confocal microscope (Carl Zeiss) equipped with 25x/0.8 OIL LD LCI PL APO or 40x/1.3 OIL DICII PL APO objectives. All quantitative imaging analyses were performed in Python.

#### Tissue segmentation

Segmentation of the organoids was performed on the DAPI channel. The channel was first blurred using a Gaussian filter and then segmented based on Otsu thresholding. The masks were later corrected manually if necessary. Embryonic explants were manually segmented.

#### Nuclei or cell segmentation and classification of cells based on their position

Nuclei were segmented in 2D using a pre-trained Stardist model ("2D_versatile_fluo") (Schmidt et al., 2018). We generated a map of the distance between each pixel within an organoid mask and the edge of the mask using the "findContours", "drawContours" and "distanceTransform" functions of the OpenCV library. We calculated the center of mass of each nucleus mask and used the distance map to classify each cell as ’external’ (distance to edge < 4.5 µm for organoids, < 10 µm for embryonic tissues) or ’internal’ (distance to edge > 4.5 µm for organoids, > 10 µm for embryonic tissues). Cells were segmented based on membrane signals (R26:mtdTomato) using Cellpose and the pre-trained model “cyto2” (Pachitariu and Stringer, 2022).

#### Hes1-emGFP and p63 intensities quantification

After nuclear segmentation based on the DAPI channels, the resulting masks were used to calculate the average signal within each mask for normalized Hes1-emGFP or p63 channels.

#### YAP nuclear to cytoplasmic ratio

A similar approach to that used for Hes1-emGFP and p63 was used to quantify the nuclear YAP signal. We then used the ’getStructuringElement’ and ’dilate’ functions of the OpenCV library to generate a ring around each nucleus, representing the cytoplasm. We then calculated the log ratio of the average YAP intensity within the nuclear mask and within the cytoplasmic ring.

#### Dll1 and Jag2 FISH quantification

After segmentation of the organoids’ global structure and of single nuclei as previously described, the smFISH was binarized and each positive pixel assigned to a nucleus based on their distance to neighboring nuclei calculated with a Voronoi diagram based on Euclidian distances.

### Explants isolation and culture

Explants were collected at two different stages of gestation. Mammary and salivary buds were collected at E13, while the urogenital sinus (UGS) and lacrimal gland were collected from E15 embryos as previously described (Berman et al., 2004; Carabaña et al., 2024; Kuony and Michon, 2017; Wang et al., 2021). Briefly, embryos were collected from the uterus of a pregnant female and stored on ice in 24-well plates. Embryos were genotyped for sex and potential transgene. Dissection of SG, LG and UGS proceeded directly, but embryos were kept in PBS at 4°C overnight before dissection of MG buds. The epidermis covering the flank and MG of the embryos was removed without enzymatic digestion.

Embryonic tissues were then placed on a cell culture insert, floating on culture medium, in 6-well plates. The embryonic culture medium contains a base of DMEM/F-12 supplemented with 1x Glutamax, 1x HEPES and 2% penicillin-streptomycin. The MG medium is supplemented with 10% fetal bovine serum (FBS) (v/v). The LG and SG medium are supplemented with 2% insulin-transferrin-selenium (ITS), while the UGS (prostate) medium also contains 10 µM dihydrotestosterone (DHT). Explant cultures were grown at 37°C with 5% CO2 and the medium was replaced every 48 hours.

### Public transcriptomic data analysis

Publicly available scRNAseq datasets of the four adult tissues and the embryonic MG, SG and UGS were retrieved from the GEO database platform. The datasets used were generated in the following studies: for adult tissues (Delcroix et al., 2023; Hauser et al., 2020; Joseph et al., 2020; Pal et al., 2017); for embryonic tissues (Carabaña et al., 2024; Farmer et al., 2017; Hauser et al., 2020; Lee et al., 2021).

## Acknowledgments

We are very grateful to Prof. Shahragim Tajbakhsh for the R26^mTmG^ reporter line and to Prof. Cedric Blanpain and Dr. Alexandra van Keymuelen for kindly providing the K5-Cre^ERT2^ mice and sample from previous publication (Centonze et al., 2020). We also wish to acknowledge to all team members, as well as our colleagues Dr. Jean-Leon Maitre, Dr. Yohanns Bellaiche, Dr. Edouard Hannezo and Prof. Michel Labouesse, for technical advice and constructive discussions. We would like to thank the Cell and Tissue Imaging Platform PICT-IBiSA (member of France-Bioimaging, ANR-10-INBS-04) of the Genetics and Developmental Biology Unit (UMR3215/U934) and the Flow Cytometry and Cell Sorting Platform at Institute Curie for their expertise, as well as the In Vivo Experimental Facility, mainly Cedrick Pauchard, Mickael Garcia and Celine Daviaud, for help in the maintenance and care of our mouse colony.

This work was funded by Paris Sciences et Lettres (PSL* Research University) (grant # C19-64-2019-228), the French National Research Agency (ANR) grant numbers ANR-21-CE13-0047 and ANR-22-CE13-0009, the Medical Research Foundation FRM "FRM Equipes" EQU201903007821, the FSER (Fondation Schlumberger pour l’éducation et la recherche) FSER20200211117, the Association for Research against Cancer (ARC) label ARCPGA2021120004232_4874, the Worldwide Cancer Research Foundation # 24-0216 and by Labex DEEP ANR-Number 11-LBX-0044 to SF.

R.J. was funded by a PhD fellowship from the French minister of research and by the FRM.

The funders had no role in study design, data collection and analysis, decision to publish, or preparation of the manuscript.

## Author Contributions

R.J. and S.F. conceived and designed the experiments. R.J., M.H. A.B. and H.M. performed all experiments. M.M.F. and J.S. contributed to experimental work and edited the manuscript. M.D. and C.F. performed mouse irradiation. R.J. developed the image analysis pipeline, performed image analysis and quantification, reanalysis of transcriptomics data, curated, and interpreted results and prepared the figures. R.J. and S.F. wrote the manuscript. S.F. provided funding, project administration and supervision. All authors reviewed and approved the manuscript.

## Materials & Correspondence

Further information and requests for resources and reagents should be directed to and will be fulfilled by the corresponding author Silvia Fre.

## Data and materials availability

All data supporting the conclusions of this study are provided in the main text or the supplementary materials.

