## Supplementary material for "Conserved signals orchestrate self-organization and symmetry breaking of bi-layered epithelia during development and regeneration"

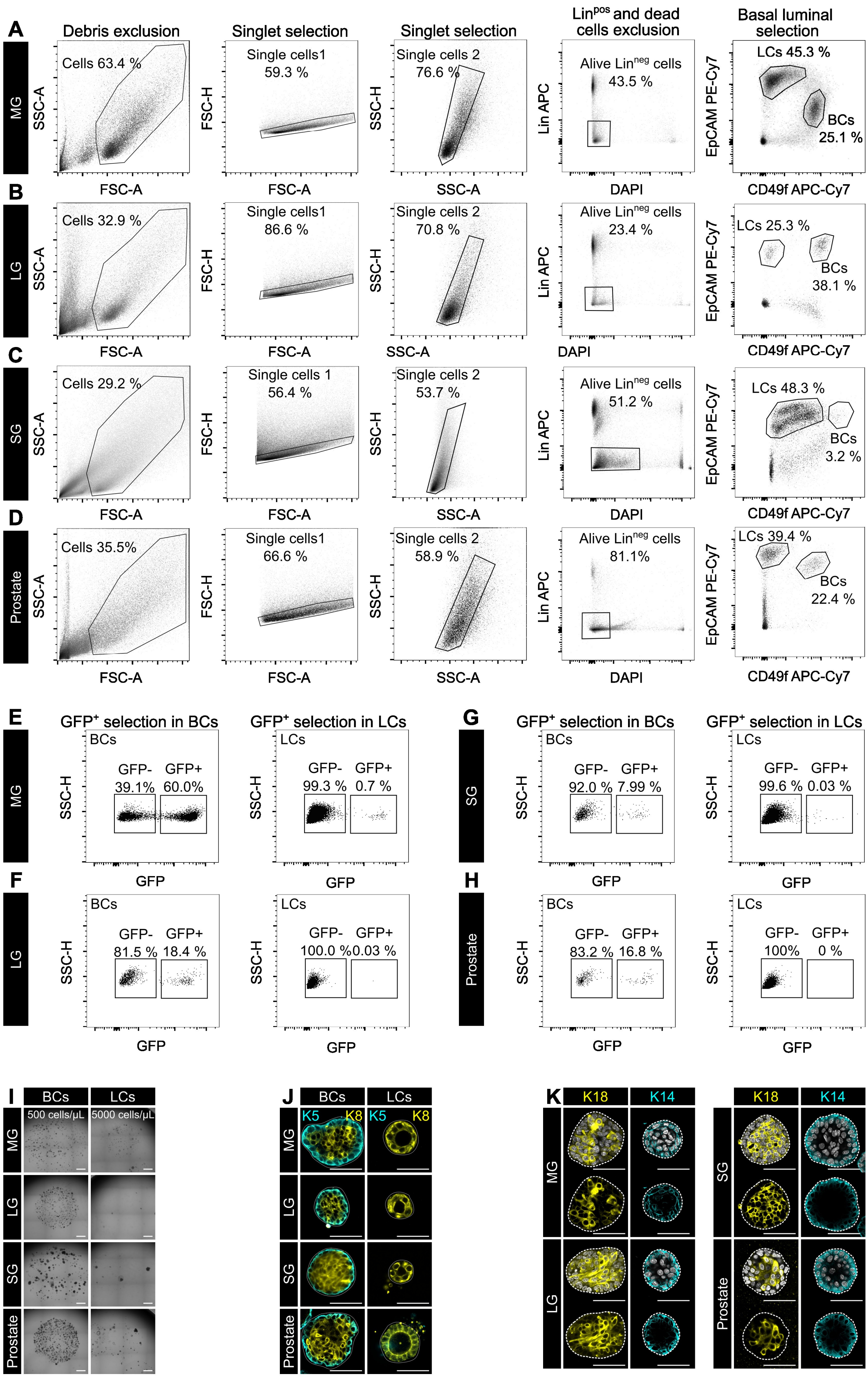

A
Debris exclusion
Singlet selection
Singlet selection
Lin^pos and dead cells exclusion
Basal luminal selection
MG
Cells 63.4 %
Single cells1 59.3 %
Single cells 2 76.6 %
Alive Lin^neg cells 43.5 %
LCs 45.3 %
BCs 25.1 %
SSC-A
FSC-H
SSC-H
Lin APC
EpCAM PE-Cy7
FSC-A
SSC-A
DAPI
CD49f APC-Cy7
B
LG
Cells 32.9 %
Single cells1 86.6 %
Single cells 2 70.8 %
Alive Lin^neg cells 23.4 %
LCs 25.3 %
BCs 38.1 %
C
SG
Cells 29.2 %
Single cells 1 56.4 %
Single cells 2 53.7 %
Alive Lin^neg cells 51.2 %
LCs 48.3 %
BCs 3.2 %
D
Prostate
Cells 35.5%
Single cells1 66.6 %
Single cells 2 58.9 %
Alive Lin^neg cells 81.1%
LCs 39.4 %
BCs 22.4 %
E
GFP+ selection in BCs
GFP+ selection in LCs
MG
BCs GFP- 39.1% GFP+ 60.0%
LCs GFP- 99.3 % GFP+ 0.7 %
G
GFP+ selection in BCs
GFP+ selection in LCs
SG
BCs GFP- 92.0 % GFP+ 7.99 %
LCs GFP- 99.6 % GFP+ 0.03 %
F
LG
BCs GFP- 81.5 % GFP+ 18.4 %
LCs GFP- 100.0 % GFP+ 0.03 %
H
Prostate
BCs GFP- 83.2 % GFP+ 16.8 %
LCs GFP- 100% GFP+ 0 %
SSC-H
GFP
I
BCs
LCs
500 cells/µL
5000 cells/µL
MG
LG
SG
Prostate
J
BCs
LCs
K5
K8
MG
LG
SG
Prostate
K
K18
K14
MG
LG
SG
Prostate

**Supplementary Figure 1. Related to Figure 1.**

**A-D**, Suspension of single cells isolated from MG (A), LG (B), SG (C) and Prostate (D) of adult *K5-Cre$^{ERT2}$:mTmG* males and females 72 h after intraperitoneal injection of TAM stained for non-epithelial markers (CD31, endothelium; CD45, immune cells; Ter119, erythroid lineage (Lin$^{pos}$ cells)) in APC, EpCAM (PE-Cy7) and CD49f (APC-Cy7) were gated to eliminate debris, doublets, Lin$^{pos}$ and dead cells, positive to DAPI incorporation. EpCAM and CD49f expression was analyzed in alive Lin$^{neg}$ cells. **E-H**, Selection of GFP$^{+}$ in BCs and LCs populations in MG (E), LG (F), SG (G) and Prostate (H). **I:** Confocal imaging of immunostaining for K14 or K18 and in organoids derived from WT BC cells. **J:** Bright field image of organoid wells seeded with BCs or LCs at define concentration in Matrigel. **K:** Confocal imaging of immunostaining for K5 and K8 of BC and LC derived organoids after 7 days in culture. Scale bar: 50 µm in **I**, J, K.

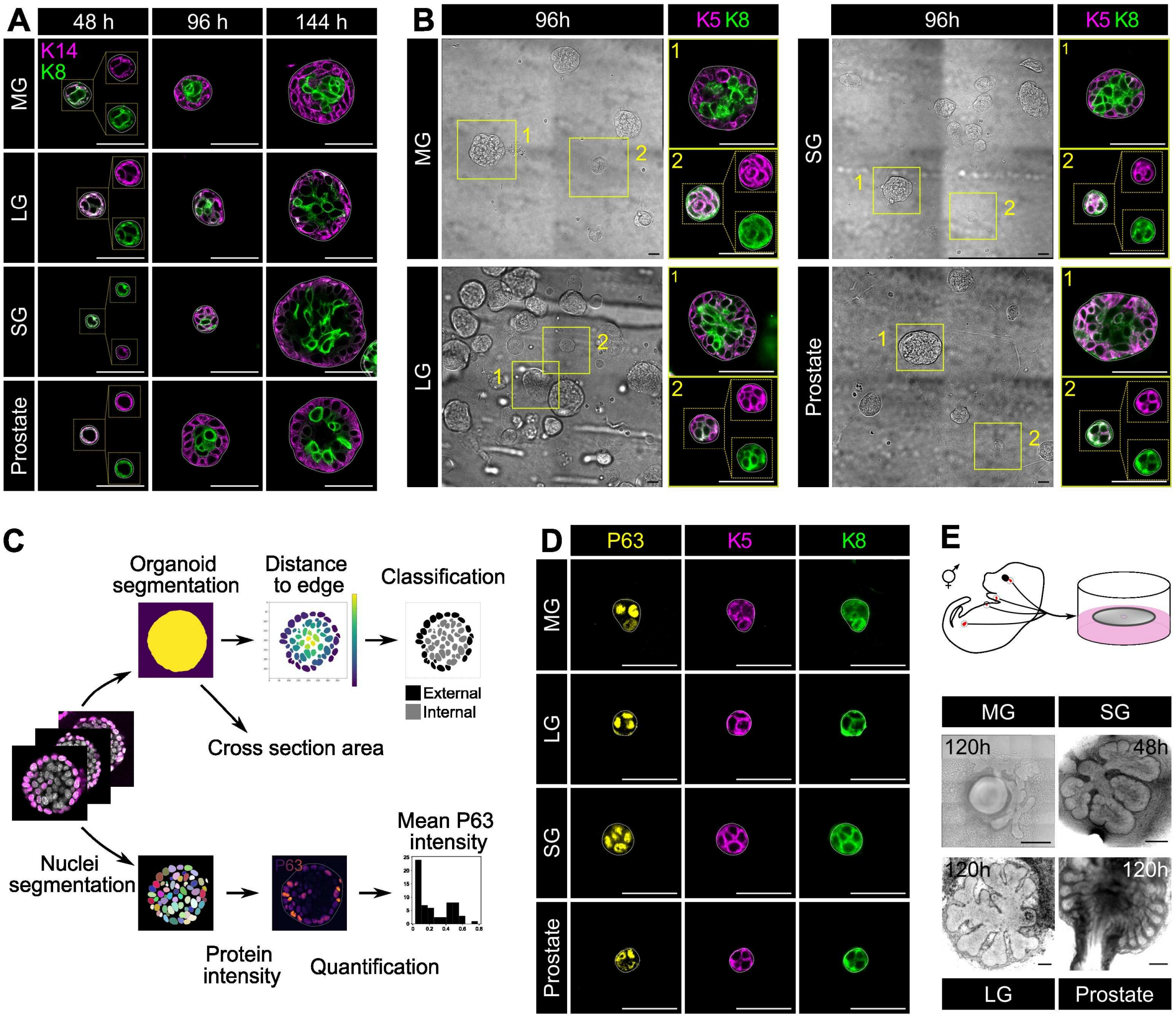

**Supplementary Figure 2. Related to Figure 2.**

**A:** Confocal imaging of immunostaining for K14 and K8 in WT organoids at different stage of growth. Scale bar: 50 µm. **B:** Bright field and confocal imaging of immunostaining for K5 and K8 in WT organoids after 96 hours in culture. Scale bar: 50 µm. **C:** Workflow of 2D quantitative image analysis of organoids. Organoids were segmented based on Otsu's thresholding and cells using Stardist on the DAPI channel. P63 intensity was quantified within each mask. See Methods section for details of image analysis. **D:** Confocal imaging of immunostaining for K5, K8 and P63 in WT organoids. Scale bar: 50 µm. **E:** Scheme of embryonic explants culture protocols and bright field imaging of explants of each tissue in culture. Scale bar: 100 µm.

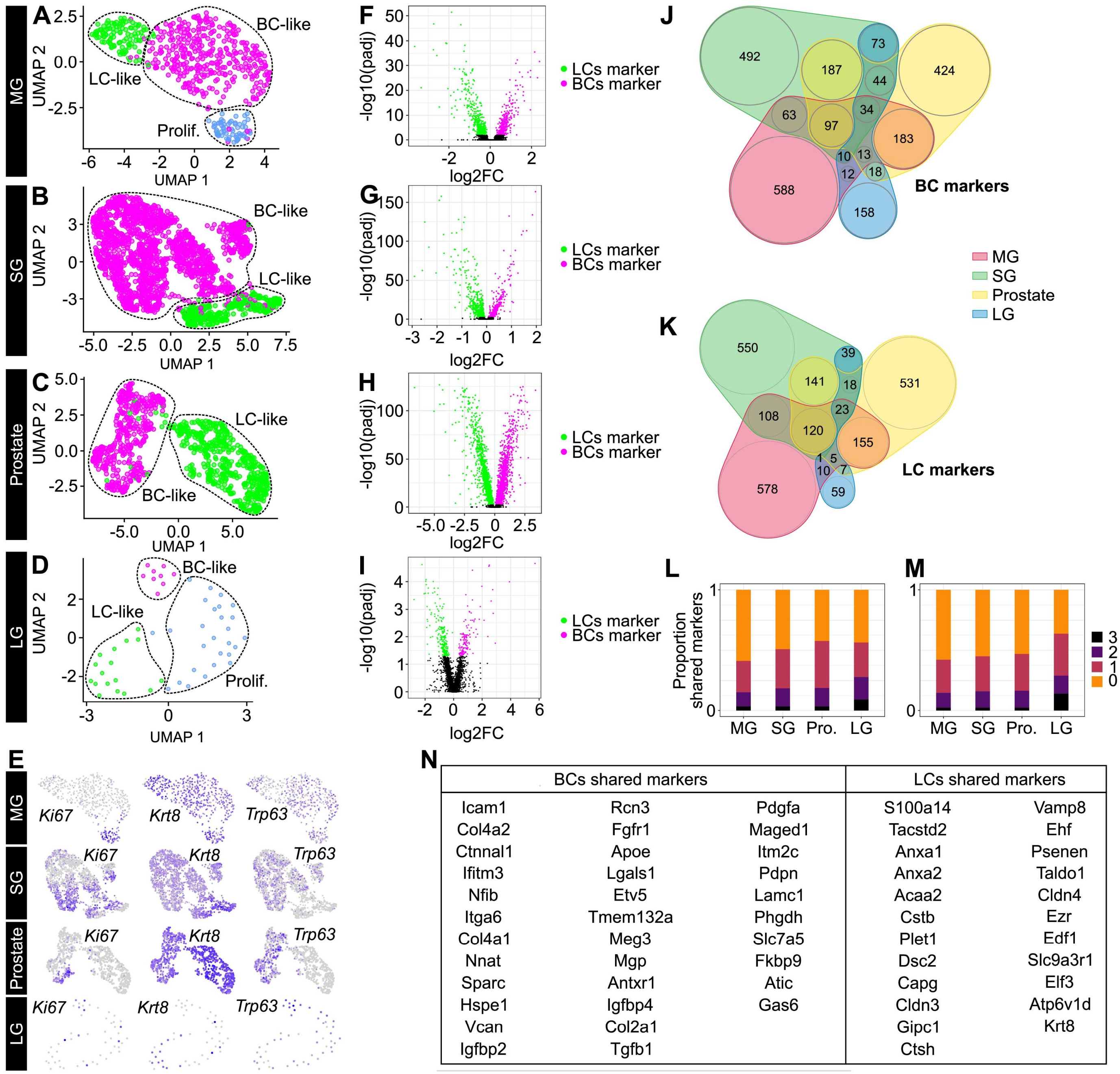

| BCs shared markers | | | LCs shared markers | |
|---|---|---|---|---|
| Icam1 | Rcn3 | Pdgfa | S100a14 | Vamp8 |
| Col4a2 | Fgfr1 | Maged1 | Tacstd2 | Ehf |
| Ctnnal1 | Apoe | Itm2c | Anxa1 | Psenen |
| Ifitm3 | Lgals1 | Pdpn | Anxa2 | Taldo1 |
| Nfib | Etv5 | Lamc1 | Acaa2 | Cldn4 |
| Itga6 | Tmem132a | Phgdh | Cstb | Ezr |
| Col4a1 | Meg3 | Slc7a5 | Plet1 | Edf1 |
| Nnat | Mgp | Fkbp9 | Dsc2 | Slc9a3r1 |
| Sparc | Antxr1 | Atic | Capg | Elf3 |
| Hspe1 | Igfbp4 | Gas6 | Cldn3 | Atp6v1d |
| Vcan | Col2a1 | | Gipc1 | Krt8 |
| Igfbp2 | Tgfb1 | | Ctsh | |

**Supplementary Figure 3. Related to Figure 3.**

**A-D**: Dimensionality reduction using UMAP of scRNA-seq data of MG E15 epithelial tissues (Carabaña et al., 2024) (**A**), SG E14.5 epithelial tissues (Hauser et al., 2020) (**B**), urogenital epithelium (UGE) E17.5 epithelial tissues (Lee et al., 2021) (**C**) and E16 LG (Farmer et al., 2017) (**D**). **E**: Expression of Ki67, Krt8 and Trp63 in the scRNA-seq presented in supplementary figure 3A-D. **F-I:** Volcano plot depicting differentially expressed genes between BCs and LCs in E15 MG (**F**), E14.5 SG (**G**), E17.5 prostate (**H**) and E16 LG (**I**). Magenta dots represent genes expressed at higher levels in BCs cells while green dots represent genes with higher expression levels in LCs/Supra-BCs (padj < 0.01). **J-K**: Venn diagram representing the shared differentially expressed genes in the BCs (**J**) or LCs (**K**) across tissues. **L-M:** Proportion of tissue specific or shared markers with 1, 2 or 3 other tissues for BCs markers (**L**) and LCs (**M**). **N:** List of BCs and LCs markers shared among the four tissues.

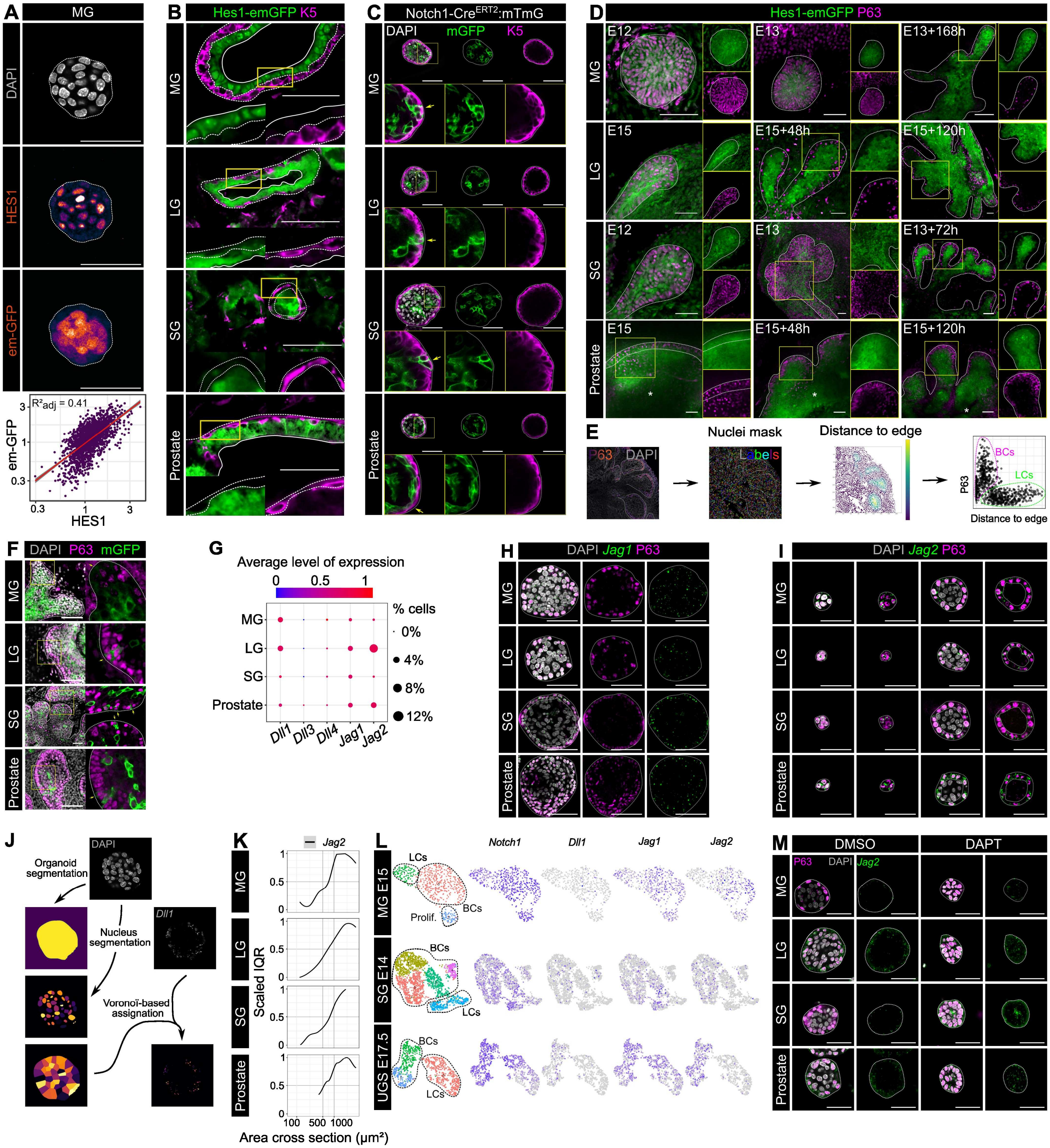

### Supplementary Figure 4. Related to Figure 4.

**A:** Confocal imaging of immunostaining for HES1 in Hes1-emGFP organoids. Scale bar: 50 µm. Relation between HES1 and Hes1-emGFP intensities in nucleus for several MG organoids. Spearman's rank correlation. **B:** Confocal imaging of immunostaining for K5 and native Hes1-emGFP in section of adult MG, LG, SG and Prostate from Hes1-emGFP animals. Scale bar: 50 µm. **C:** Confocal imaging of immunostaining for K5 and native mGFP in *Notch1-Cre$^{ERT2}$:mTmG* organoids after 7 days in culture. Organoids were incubated with 0.5 µM 4-OHT for 24h before fixation. Scale bar: 50 µm. **D:** Confocal imaging of immunostaining for P63 and native Hes1-emGFP explants at different stages of growth. Scale bar: 50 µm. **E:** Workflow of 2D quantitative image analysis of explants. The epithelium was manually segmented, while nuclei were segmented using Stardist. The distance between each nucleus and epithelial limit is calculated by 2D projection and markers intensities were then quantified within each mask. See Methods section for details of image analysis. **F:** Confocal imaging of immunostaining for P63 and native mGFP in Notch1-Cre$^{ERT2}$:mTmG explants. Explants were incubated with 0.5 µM 4-OHT for 24h before fixation (MG: E13+168h, SG: E13+72h, LG: E15+120h, Prostate: E15+120h). Scale bar: 50 µm. **G:** Dot plot representing the expression of Notch ligands in adult MG, LG, SG and Prostate tissues. Data were extracted from published scRNA seq data of adult tissues (See Method section). Size of dots corresponds to the percentage of cells expressing the markers in the epithelial cluster positive for EpCAM. The level of expression in expressing cells is color-coded. **H:** Confocal imaging of smFISH for *Jag1* and immunostaining for P63 in WT organoids after 7 days in culture. Scale bar: 50 µm. **I:** Confocal imaging of smFISH for *Jag2* and immunostaining for P63 in WT organoids at different stages of growth. Scale bar: 50 µm. **J:** Workflow of 2D quantitative image analysis of smFISH signal in organoids. Organoids are segmented based on Otsu's thresholding, nuclei segmented using Stardist and smFISH signal binarized. Each binarized pixel is assign to the closest nucleus based on a Voronoi diagram. See Methods section for details of image analysis. **K:** Interquartile range (Q3-Q1) of the *Jag2* signal measured in each cell of a center optical cross section of organoids of different size (n=3 independent cell sorting experiments). See Methods section. **L:** Expression of Notch ligands and receptors in epithelial embryonic MG (Carabaña et al., 2024), SG (Hauser et al., 2020) and Prostate tissue (Lee et al., 2021). **M:** Confocal imaging of smFISH for *Jag2* and immunostaining for P63 in WT organoids treated with DMSO or 50 µM DAPT for 7 days. Scale bar: 50 µm.

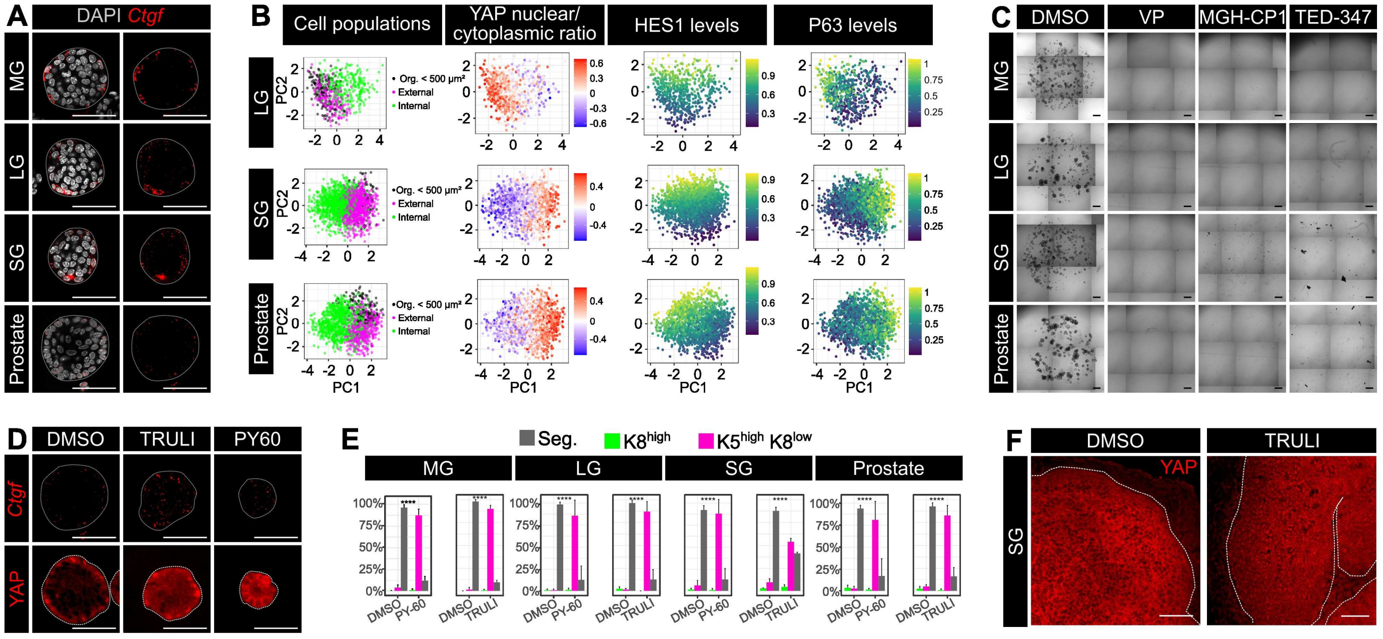

### Supplementary Figure 5. Related to Figure 5.

**A:** Confocal imaging of smFISH for *Ctgf* (red dots) in WT organoids after 7 days in culture. Scale bar: 50 µm. **B:** Principal component analysis (PCA) of cells stained for YAP and P63 in Hes1-emGFP LG, SG and Prostate derived organoids of different size (n=3 independent cell sorting experiments). See Methods section. **C:** Bright field image of BCs derived organoids after 7 days in culture with DMSO or YAP inhibitors, Verteporfin (VP) (2 µM), MGH-CP1 (10 µM) and TED-347 (10 µM). Scale bar: 500 µm. **D:** Confocal imaging of smFISH for *Ctgf* or immunostaining for YAP in WT prostate organoids treated with DMSO, 20 µM TRULI or 10 µM PY-60 for 7 days in culture. Scale bar: 50 µm. **E:** Quantification of the percentage of organoids composed of $K8^{high}$ cells only, $K5^{high}K8^{low}$ only or a mix of $K5^{high}K8^{low}$ and $K5^{low}K8^{high}$ cells. Data are displayed as Mean + sd (n=3 independent cell sorting experiments). Chi square test. **F:** Confocal imaging of immunostaining for YAP in WT SG explants treated for 72 h with DMSO or 20 µM TRULI. The dotted line delineates the epithelium limit Scale bar: 50 µm.

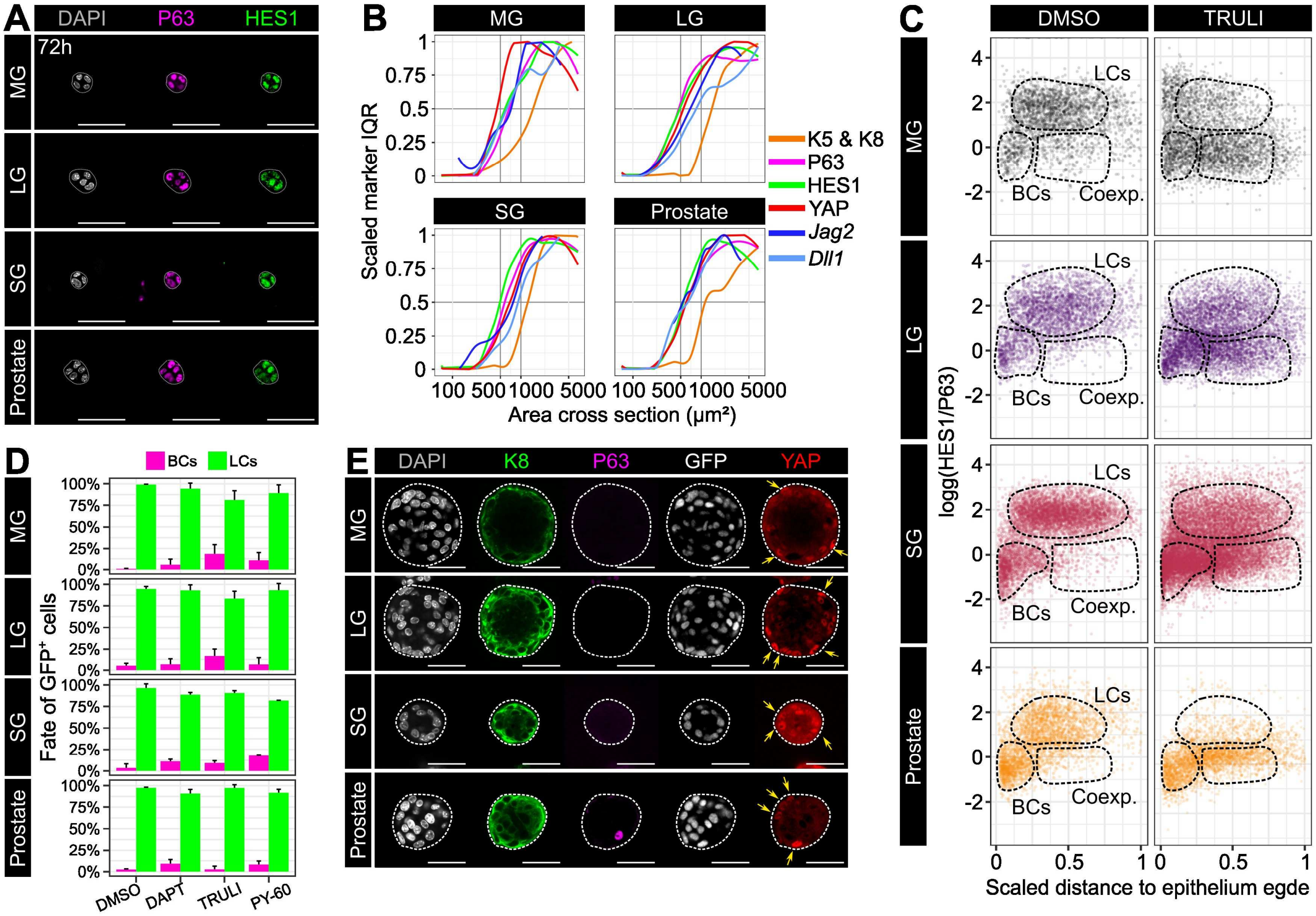

**Supplementary Figure 6. Related to Figure 6.**

**A:** Confocal imaging of immunostaining for p63 and HES1 in WT organoids after 72 hours in culture. Scale bar: 50 µm. **B:** Scaled Interquartile range of K5 & K8, P63, HES1 YAP, *Dll1* and *Jag2* in organoids of different size (n=3 independent cell sorting experiments). **C:** HES1 and p63 log ratio in cells from explants cells treated with DMSO or 20 µM TRULI for the whole duration of the culture in relation with their normalized distance to the epithelium limit (n=3 biological replicates). **D:** Fate of N1ICD-nGFP$^{+}$ cells *in K5-Cre$^{ERT2}$:N1ICD-nGFP* organoids treated for 7 days with DMSO, 50 µM DAPT, 20 µM TRULI or 10 µM PY60. Data are displayed as Mean + sd (n=3 independent cell sorting experiments). **E:** Confocal imaging of immunostaining for p63, K8, YAP and N1ICD-nGFP in *K5-Cre$^{ERT2}$:N1ICD-nGFP* organoids with high percentage of recombination after 7 days in culture with initial treatment with 0.5 µM 4-OHT for the first 48 hours in culture. Scale bar: 50 µm.

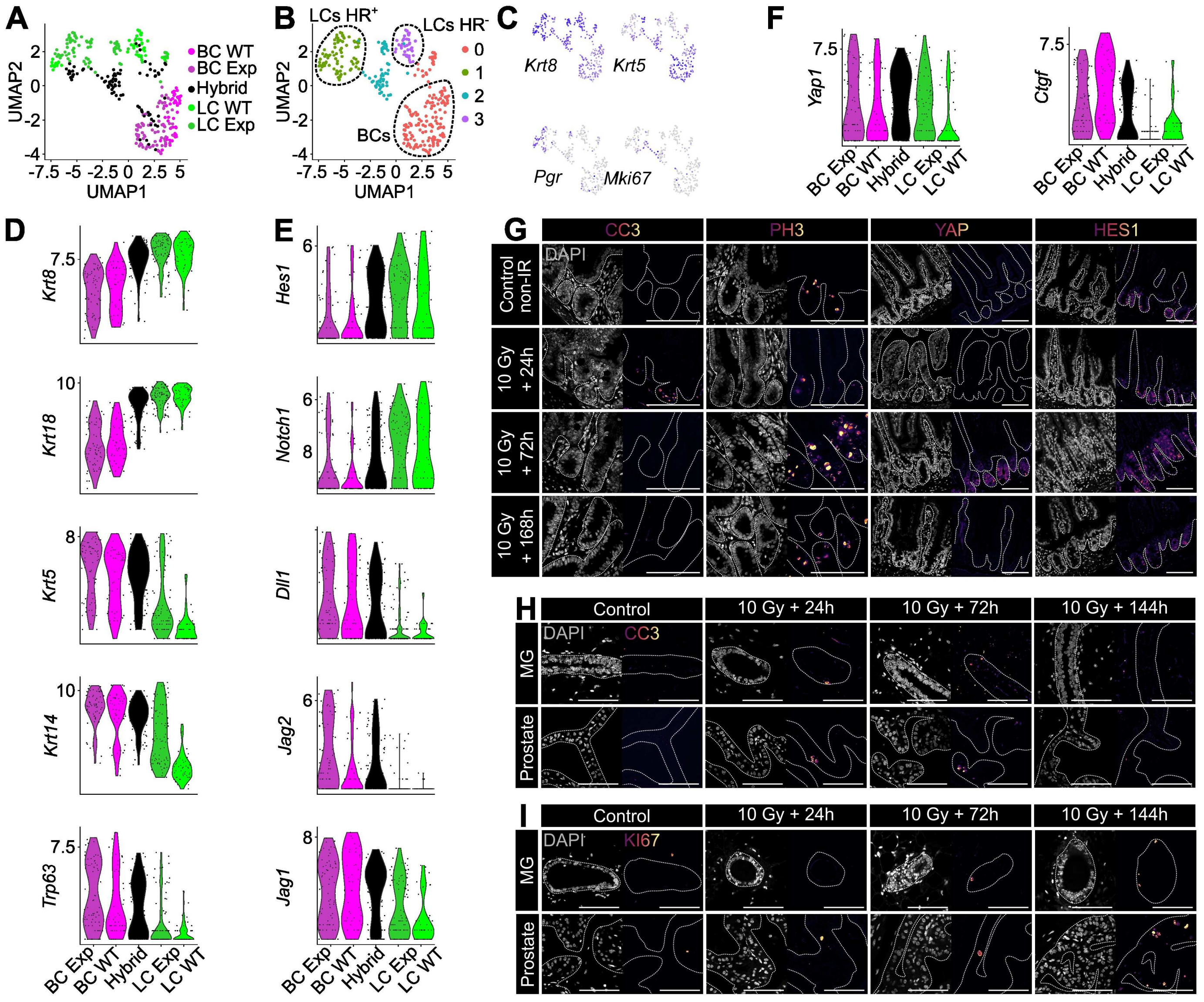

**Supplementary Figure 7. Related to Figure 7.**

**A-B:** Dimensionality reduction using UMAP of scRNA-seq data of MG epithelial tissues cells sequenced 1 week post ablation and after labelling of BCs (Centonze et al., 2020). Cell population identified during sorting (**A**) or by gene signature (**B**) are color-coded. **C:** Expression of the indicated differentiation markers projected onto the UMAP dimensionality reduction displayed in A and B. **D-F:** Violin plot of expression level for indicated markers of fate (**D**), Notch (**E**) or Hippo (**F**) pathways components in the different FACS related population. **G:** Confocal imaging of immunostaining for cleaved Caspase 3 (CC3), Phospho Histone 3 (PH3), YAP and HES1 in WT adult tissues before or at different time points after irradiation. Scale bar: 50 µm. **H:** Confocal imaging of immunostaining for cleaved Caspase 3 (CC3) in WT adult tissues before or at different time points after irradiation. Scale bar: 50 µm. **I:** Confocal imaging of immunostaining for KI67 in WT adult tissues before or at different time points after irradiation. Scale bar: 50 µm.

### Key Resource Table 1. Reagents and materials

| Reagent | Source | Reference | Concentration | Organoid medium |
|---|---|---|---|---|
| **Chemicals, peptides and recombinant proteins** | | | | |
| DMEM/F12 | Gibco | 21331020 | 1x | MG, LG, SG, Pro |
| Glutamax | Thermo Fisher Scientific | 35050038 | 1x | MG, LG, SG, Pro |
| HEPES (1M) | Gibco | 15630080 | 1x | MG, LG, SG, Pro |
| B27 supplement | Thermo Fisher Scientific | 12587-010 | 2x | MG, LG, SG, Pro |
| N2 supplement | Thermo Fisher Scientific | 17502048 | 1x | MG |
| Penicillin/Streptomycin | Thermo Fisher Scientific | 15140122 | 2% | MG, LG, SG, Pro |
| FGF STAB | Enantis | CAT. NO.: FGF2STAB01000 | 50ng/mL | MG, LG, SG, Pro |
| mNoggin | PeproTech | 250-38 | 100 ng/mL | MG, LG, SG, Pro |
| mRspo1 | PeproTech | 3474-RS | 100 ng/mL | MG, LG, SG, Pro |
| N-acetylcysteine | Medchemexpress | HY-B0215 | 1.25 mM | SG, LG, Pro |
| Neuregulin | R&D Systems | 5898-NR-050 | 100ng/mL | MG |
| Wnt3a | R&D Systems | 1324-WNP-010/CF | 5 ng/µL | LG, SG |
| mEGF | Thermo Fisher Scientific | 315-09 | 2 ng/µL | LG, SG, Pro |
| A8301 | TOCRIS | 2939 | 10 µM | Pro |
| Dihydro-testosterone (DHT) | Sigma-Aldrich | A8380 | 1 nM (organoids), 10 µM (explants) | Pro |
| Y27632 | Medchemexpress | HY-10071 | 10 µM | MG, LG, SG, Pro |
| Collagenase A | Roche | 10103586001 | | |
| DNAse I | Sigma-Aldrich | D4527 | | |
| Dispase II | Roche | 10888700 | | |
| Sunflower Oil | Sigma-Aldrich | S5007 | | |
| Tamoxifen | MP Biomedicals | 156738 | 2mg/10g of mouse | |
| 4-OHT | Medchemexpress | HY-16950 | | |
| Cell Recovery Solution | Corning | 354253 | 1x | |
| EDTA | MilliporeSigma | E6765 | | |
| CO2 independent | Gibco | 18045054 | 1x | |
| Fetal bovine serum (FBS) | Thermo Fisher Scientific | 10500064 | 10% | |
| Insulin Transferrin Selenium (ITS) | Gibco | 41400045 | 2% | |
| Paraformaldehyde (PFA) | Electron Microscopy Sciences | 15710 | 4% | |
| Aqua poly/mount | Tebu Bio | 18606-5 | | |
| Glycerol | Euromedex | 15710 | | |
| Triton X-100 | Euromedex | 2000C | | |
| Sucrose | Sigma | S0389 | | |

| | | |
|---|---|---|
| Tissue-Tek O.C.T. | Sakura | 4583 |
| DAPT | Medchemexpress | Cat. No.: HY-13027 |
| TRULI | Medchemexpress | HY-138489 |
| PY60 | Medchemexpress | HY-141644 |
| Verteporfin | Medchemexpress | HY-B0146 |
| RNAscope Multiplex Fluorescent Detection Kit v2 kit | ACD | 32310 |
| RNAscope H2O2 and protease reagents | ACD | 322381 |
| RNAscope Target Retrieval Reagent | ACD | 322000 |
| RNAscope TSA buffer pack | ACD | 322810 |
| RNAscope Probe Diluent | ACD | 300041 |
| TSA PLUS FLUORESCEIN | Akoya biosciences | NEL741001KT |
| TSA PLUS CYANINE 3 | Akoya biosciences | NEL744001KT |
| TSA PLUS CYANINE 5 | Akoya biosciences | NEL705A001KT |
| RNAscope® 3-plex positive & Negative Control Probe | ACD | 320871 |
| RNAscope® Probe-Mm-Notch1 | ACD | 404641 |
| RNAscope® Probe-Mm-Jag1 | ACD | 412831 |
| RNAscope® Probe-Mm-Jag2 | ACD | 417511 |
| RNAscope® Probe-Mm-Dll1 | ACD | 425071 |
| RNAscope® Probe-Mm-Ctgf | ACD | 314541 |

**Key Resource Table 2. Antibodies**

| Reagent | Source | Reference | Dilution |
|---|---|---|---|
| Antibodies | | | |
| Rabbit anti-K5 | Covance | Cat# PRB-160P-100; RRID: AB_291581 | 1:300 |
| Rat anti-K8 | Developmental Studies Hybridoma Bank, University of Iowa | Cat# TROMA-I: RRID: AB_531826 | 1:300 |
| Mouse anti-p63 | Abcam | Cat# ab735; RRID: AB_305870 | 1:200 |
| Chicken anti-K14 | Biolgend | # 906004 | 1:300 |

| | | | |
|---|---|---|---|
| Rat anti-K18 | | Kemler et al. 1981 | 1:200 |
| Rabbit anti-CC3 | Cell Signaling Technology | Cat# 9664; RRID:AB_2070042 | 1:200 |
| Rabbit anti-Ki67 | Thermo Fisher Scientific | Cat# MA5-14520; RRID:AB_10979488 | 1:300 |
| Rabbit anti-ERα | Agilent-Dako | Cat# M7047, RRID: AB_2101946 | 1:100 |
| Rabbit anti-PR | SantaCruz | sc-7208; RRID: AB_2164331 | 1:100 |
| Rabbit anti-PAX6 | Developmental Studies Hybridoma Bank | AB2237 | 1:300 |
| Rabbit anti-MIST1 | Abcam | ab187978 | 1:50 |
| Rabbit anti-AR | Proteintech | 22089-1-AP | 1:100 |
| Rabbit anti-HES1 | Cell Signaling technology | #11988 | 1:100 |
| Human anti-GFP | | Recombinant anti GFP antibody | 1:200 |
| Rabbit anti-YAP | Cell Signaling Technology | Cat# 14074 | 1:300 |
| Goat anti-rabbit AlexaFluor-coupled to different fluorochromes (Cy3, Cy5, A488, A405) | Invitrogen-Thermo Fisher Scientific | Cat# A10520; RRID: AB_2534029 Cat#A10523; RRID: AB_2534032, Cat# A-11034; RRID: AB_2576217 Cat# A-31556; RRID: AB_221605 | 1:500 |
| Goat anti-rat AlexaFluor-coupled to different fluorochromes (Cy3, Cy5, A488) | Invitrogen-Thermo Fisher Scientific | Cat# A10522; RRID: AB_2534031, Cat#A10525; RRID: AB_2534034, Cat# A-11006; RRID: AB_2534074 | 1:500 |
| Goat anti-mouse AlexaFluor-coupled to different fluorochromes (Cy3, Cy5, A488) | Invitrogen-Thermo Fisher Scientific | Cat# A10521; RRID: AB_2534030, Cat# A10524; RRID: AB_2534033 Cat# A-11001; RRID: AB_2534069 | 1:500 |
| EpCAM PE/Cy7 | Biolegend | Cat# 118216; RRID: AB_1236471 | 1:100 |
| CD49f APC/Cy7 | Biolegend | Cat# 313628; RRID: AB_2616784 | 1:100 |
| CD31 APC | Biolegend | Cat# 102510; RRID: AB_312905 | 1:100 |
| Ter119 APC | Biolegend | Cat# 116212; RRID: AB_313713 | 1:100 |
| CD45 APC | Biolegend | Cat# 103112, RRID: AB_312977 | 1:100 |

**Key Resource Table 3. Animals**

| Strain | Source | Reference | Common name |
|---|---|---|---|
| JAX stock 008463 | RRID:IMSR_JAX:008463 | (Ventura et al., 2007) | *R26-CreERT2* |
| JAX stock 029155 | RRID:IMSR_JAX:029155 | (Van Keymeulen et al., 2011) | *K5-CreERT2* |
| | | (Fre et al., 2011) | *N1-CreERT2* |
| | | (Murtaugh et al., 2003) | *R26-N1ICD* |
| | | (Fre et al., 2011) | *HES1-emGFP* |
| JAX stock: 037456 | RRID:IMSR_JAX:037456 | (Muzumdar et al., 2007) | *R26-mTmG* |
